# Engineered cell-like vesicles instruct keratocyte differentiation for corneal biofabrication and regeneration

**DOI:** 10.64898/2026.08.06.743353

**Authors:** Alexandre Taoum, Ole Thaden, Antonette Jessica Arunkumar, Philipp Scheulen, Christopher Robin Wood, Andrea Frank, Meng Wang, Friederike Dehli, Daniela F. Duarte Campos

## Abstract

Precise control of growth factor delivery remains a challenge for directing stem cell differentiation in three-dimensional (3D) engineered tissues. In this study, engineered cell-like vesicles are used as programmable microenvironments to enable sustained and localized delivery of growth factors within visible-light-crosslinked GelMA hydrogels. Giant unilamellar vesicles (GUV) loaded with FGF-2 and TGF-β3 were incorporated into bioinks with BM-MSC to drive keratocyte differentiation without repeated soluble growth factor supplementation. ELISA measurements confirmed the removal of non-encapsulated growth factors and the release of the vesicle cargo following induced vesicle rupture. Fluorescence monitoring showed a progressive reduction in detectable FGF-2- and TGF-β3-loaded GUV during culture, while droplet-scale analysis demonstrated the co-deposition of cells and vesicles after printing. After 14 days of differentiation, differentiated cells expressed ALDH1A1, ALDH3A1, lumican, keratocan, and collagen I without induction of α-SMA. Interestingly, keratocyte-associated differentiation was retained after drop-on-demand bioprinting, confirmed by qPCR analysis. These findings establish growth factor-loaded vesicles as bioprintable instructive niches capable of supporting localized keratocyte differentiation within 3D corneal constructs.

## Introduction

The ability to direct stem cell differentiation within engineered tissues depends on the establishment of a microenvironment in which biochemical signals are delivered at an appropriate concentration, location, and time[1–5]. In conventional three-dimensional (3D) culture, differentiation-inducing factors are generally supplied through the culture medium and refreshed repeatedly throughout differentiation period[6, 7]. Although this approach is suitable for *in vitro* studies, soluble supplementation provides limited spatial control, remains dependent on repeated external intervention, and can be affected by dilution, diffusion, and loss of growth factor activity during prolonged culture[5, 8–10]. These limitations become particularly relevant when differentiation is intended to occur directly within a bioprinted construct.

To tackle these limitations, a range of delivery strategies has been explored, including extracellular vesicles[11–13] and nanoparticle-based systems[14–16]. These approaches have demonstrated the capacity to modulate stem cell behavior by transporting bioactive molecules and activating intracellular signaling pathways[17–19]. However, their ability to recapitulate complex and sustained microenvironmental signaling remains limited, particularly within 3D engineered constructs[11, 20, 21]. In this context, while extracellular vesicles and nanoparticle-based systems have been already investigated, the use of synthetic, cell-sized vesicles as programmable microenvironments to direct mesenchymal stromal cell differentiation within 3D bioprinted constructs remains, to our knowledge, unexplored.

Synthetic cell-like vesicles offer a unique opportunity to address this gap by enabling the engineering of localized and tunable signaling niches. Giant unilamellar vesicles (GUV), due to their cell-scale dimensions and compartmentalization capacity, can encapsulate defined combinations of biomolecules and release them in a controlled manner[22–24]. In contrast to conventional delivery systems, such vesicles can function as spatially confined sources of morphogens, thereby providing sustained and gradient-dependent biochemical cues within complex 3D environments[10, 25–27].

This concept could be particularly relevant in the context of corneal tissue engineering, where the induction of a keratocyte phenotype from mesenchymal stromal cells (MSC) is dependent on sustained exposure to specific growth factors, including fibroblast growth factor-2 (FGF-2) and transforming growth factor-β3 (TGF-β3)[28–31]. Current differentiation protocols in 3D constructs require prolonged culture periods and repeated supplementation to compensate for growth factor instability at physiological temperature[32, 33]. For example, such requirements are incompatible with *in situ* bioprinting strategies[34], where repeated and self-delivery of soluble growth factors would be required to enable differentiation.

To make this strategy compatible with drop-on-demand bioprinting, both biological and processing requirements must be fulfilled. Growth factors must remain encapsulated after purification[35–37], vesicles must be detectable within the hydrogel over the culture period[38, 39], and the complete cell- and vesicle-containing formulation must remain cytocompatible[40–42] without altering the mechanical properties of the hydrogel[43, 44]. In addition, bioinks including cells and vesicles must be printable with reproducible spatial distributions[22, 45, 46]. Finally, vesicle-mediated signaling must remain sufficient to induce a keratocyte-associated phenotype after exposure to the printing process.

In this study, FGF-2-loaded and TGF-β3-loaded vesicles were incorporated with BM-MSC into visible-light-crosslinked riboflavin-arginine gelatin methacryloyl (RA-GelMA). Vesicle purification and growth factor encapsulation were assessed by enzyme-linked immunosorbent assay (ELISA), while fluorescence imaging and quantitative vesicle counting were used to examine vesicles persistence during culture. The storage modulus of the complete BM-MSC- and vesicle-containing formulation, long-term cell viability, and the radial distribution of cells and vesicles following drop-on-demand printing were evaluated. Keratocyte differentiation was assessed in cast and printed constructs by immunofluorescence and qRT-PCR, using ALDH1A1, ALDH3A1, lumican, keratocan, and collagen I as keratocyte-associated markers[47–50] and α-SMA as an indicator of myofibroblastic activation[51, 52].

This work establishes synthetic cell-like vesicles as a platform for engineering instructive microenvironments in 3D biofabrication, providing a strategy to program cell differentiation within bioprinted tissues. Beyond corneal regeneration, this approach may be broadly applicable to the development of other functional engineered tissues.

## Materials and Methods

### Synthetic growth factor filled vesicle preparation

Lipids 1,2-di(9Z-octadecenoyl)-*sn*-glycero-3-phosphocholine (DOPC), 1,2-dioleoyl-*sn*-glycero-3-phosphoethanolamine-*N*-[methoxy(polyethylene glycol)-2000] (ammonium salt) (PEG(2000)-PE), 1,2-dioleoyl-*sn*-glycero-3-phosphoethanolamine-*N*-(lissamine rhodamine B sulfonyl) (ammonium salt) (Liss Rhod-PE) and 1,2-dioleoyl-sn-glycero-3-phosphoethanolamine-N-(Cyanine 5) (Cy5-PE) were purchased from Avanti Research, Alabaster, USA. All lipids were stored in chloroform (288306, Sigma-Aldrich, St. Louis, USA) at −20°C.

PEGylated growth factor-loaded giant unilamellar vesicles (GUV) were produced by electroformation as previously described by Thaden et *al*.[22]. Briefly, lipid mixtures consisted of 94 mol % 1,2-dioleoyl-sn-glycero-3-phosphocholine (DOPC), 5 mol % PEG(2000)-PE, and 1 mol % fluorescent lipid, either Lissamine Rhodamine-PE (for FGF-2-loaded GUV) or Cy5-PE (for TGF-β3-loaded GUV). Lipids were dissolved in chloroform (3 mM), and 50 µl of the solution was spread onto indium tin oxide (ITO)-coated glass slides and allowed to dry under a fume hood for 30 min. An electroformation chamber was assembled using an 18 mm rubber spacer, and 270 µl of growth factor solution containing 300 mM sucrose (S0389, Sigma-Aldrich) supplemented with either 1 µg mL⁻¹ FGF-2 or 0.1 µg mL⁻¹ TGF-β3 was introduced. The chamber was sealed with a second ITO-coated slide and subjected to an alternating electric field (3 Vpp, 5 Hz) at 37 °C for 128 min to induce vesicle formation.

Following electroformation, vesicle suspensions were collected, washed to remove non-encapsulated growth factors, and concentrated. Briefly, samples were diluted 1:20 in 300 mM glucose solution, centrifuged at 200 × g for 10 min, and the supernatant was partially removed to a final volume of 50 µl. This washing step was repeated three times. Vesicle concentration was determined by fluorescence imaging using a counting chamber, and cell-sized vesicles (diameter > 8 µm) were quantified using GUVdetector (University Cologne)[53]. Vesicles were used on the day of preparation.

For co-encapsulation, both growth factors were incorporated during a single electroformation process following the same protocol.

### RA-GelMA preparation

RA-GelMA preparation was conducted following the protocol of Dehli et *al.*[54]. Briefly, 25.75 g of gelatin type B (G9391, 225 Bloom, Sigma Aldrich, Germany) was dissolved in 250 mL deionized water at a temperature of 40 °C. The temperature was lowered to 37 °C and 13 mL of methacrylic anhydride was added. The pH was adjusted to 7.3 by continuous addition of 4 M NaOH *via* an automatic titrator (T50, Mettler Toledo, Germany). The reaction time was 5 h. Afterward, the reaction mixture was filtered and the crude product was stored at 8 °C for two days. The synthesized GelMA was purified by dialysis against deionized water using a 12-14 kDa MW cutoff dialysis membrane (Sigma Aldrich). The dialysis was done for 5 days, changing the water twice a day. ^1^H-NMR (300 MHz) in D_2_O was used to determine the degree (DM) of methacryloylation using TMSP as an internal standard. GelMA with a DM of 0.77 mmol g^−1^ was used for all experimental work.

### Cell culture

Human bone marrow-derived mesenchymal stromal cells (BM-MSC) from three donors (male, 63 years; female, 62 and 72 years; all Caucasian) were obtained from PromoCell (C-12974) and pooled prior to expansion. Cells were subcultured in Mesenchymal Stromal Cell Growth Medium 2 (C-28009, PromoCell) supplemented with 1% penicillin-streptomycin (P4333, Sigma-Aldrich).

### Preparation and processing of RA-GelMA with cells and vesicles

Cells were detached using Accutase (A6964, Sigma-Aldrich) and encapsulated at a concentration of 2 × 10⁶ cells mL⁻¹ in 30 wt% riboflavin-arginine gelatin methacryloyl (RA-GelMA) dissolved in culture medium. The precursor solution was supplemented with 190 µmol l⁻¹ riboflavin (CDS025203, Sigma-Aldrich) and 80 mmol l⁻¹ L-arginine monohydrochloride (A5131, Sigma-Aldrich).

For conditions using separately prepared vesicles, FGF-2-loaded vesicles were incorporated at 2 × 10⁶ vesicles mL⁻¹ and TGF-β3-loaded vesicles at 1 × 10⁶ vesicles mL⁻¹. For co-encapsulation, both growth factors were loaded within the same vesicle population. To achieve an equivalent effective dose to the separate condition, loading ratio of TGF-β3 was adapted to 1:2, with a final total vesicle concentration of 2 × 10⁶ vesicles mL⁻¹. The resulting hydrogel precursor solution was then cast into cylindrical molds (2 mm height, 4 mm diameter) prior to crosslinking.

Photocrosslinking was performed for 10 min using visible blue light (455 nm LED, Thorlabs) at an intensity of 4.3 mW cm⁻². Following crosslinking, constructs were transferred into 1 mL of culture medium and incubated for 1 h to allow equilibration. The medium was then replaced, and samples were maintained in 1 mL of their respective culture conditions.

The stability of encapsulated GUV within RA-GelMA was assessed by fluorescence imaging. Z-stack images were acquired at defined time points of the differentiation process (day 0, 7, 14) using the EVOS (M5000, Thermo Fisher Scientific) fluorescence microscope (20X magnification; 50 slices, 2.5 µm step size). Image post-processing, including background subtraction, was performed using IMARIS (Version 10.2.0, Oxford Instruments), the GUV segmented and counted. The same 3D constructs were imaged over time to enable consistent evaluation of vesicle integrity within the hydrogel matrix.

### 3D culture and differentiation

Differentiation of control samples was performed according to the protocol described by Taoum et al. (2025). Briefly, constructs were cultured for 7 days post-crosslinking prior to induction, after which they were incubated for 14 days in keratocyte differentiation (KD) medium. The KD medium consisted of DMEM/F12 supplemented with 1% MEM vitamin solution, 1% non-essential amino acids, 1% insulin-transferrin-selenium (41400045, Gibco), 1 mM L-ascorbate 2-phosphate, 20 ng mL⁻¹ fibroblast growth factor-2 (CB-1102021, Pan-Biotech), 0.1 ng mL⁻¹ transforming growth factor-β3 (78131, STEMCELL Technologies), 1% amphotericin B, and 1% penicillin-streptomycin. The medium was refreshed every 2 days.

For vesicle-driven differentiation, induction was initiated 3 days post-crosslinking. In this condition, the delivery of fibroblast growth factor-2 and transforming growth factor-β3 was achieved through sustained release from encapsulated vesicles. Constructs were therefore cultured in growth factor-free KD medium for 14 days, with medium replacement every 2 days.

### Bioprinting of RA-GelMA bioinks

Cell-laden bioinks were prepared as described above and printed using a drop-on-demand bioprinter (SuperFill, custom design, Black Drop Biodrucker GmbH, Germany) equipped with an electromagnetic microvalve (nozzle diameter 600 µm, Fritz Gyger AG, Switzerland). Printing was performed using a pressurized air supply at a constant pressure of 1 bar.

Three experimental conditions were investigated following printing: (i) no differentiation, (ii) standard differentiation with soluble growth factor supplementation, and (iii) vesicle-mediated differentiation. For all conditions, constructs were printed directly into the molds and crosslinked immediately after deposition under visible light, as described above.

Printed constructs were maintained under standard culture conditions, and differentiation protocols were initiated according to the respective experimental group. In the normal differentiation condition, constructs were cultured in keratocyte differentiation medium supplemented with soluble growth factors, whereas in the vesicle-mediated condition, growth factor delivery was achieved through encapsulated vesicles incorporated within the bioink.

### Bioink printability and distribution analysis

Bioinks containing 0, 10, 20, 30, and 40 wt % RA-GelMA were printed using combinations of printing pressures (0.25, 0.5, 1.0, and 1.5 bar) and valve opening times (250, 500, 750, and 1000 µs). Each printing condition was evaluated in triplicate and considered printable only when all three independent trials produced reproducible droplet generation. Conditions resulting in nozzle clogging or the absence of droplet formation were classified as non-printable.

The spatial distribution of both BM-MSC and vesicles with a concentration of 1 ×10⁶ cells mL⁻¹ and 1 ×10⁶ vesicles mL⁻¹ within printed 30 wt % RA-GelMA droplets was evaluated following drop-on-demand bioprinting. For each independent experiment, 10 individual droplets were analyzed, with three technical replicates included (T1 to T3).

Printed droplets were imaged by brightfield and fluorescence microscopy. Brightfield images were used to define the droplet boundary, while fluorescence images were acquired to visualize cell nuclei and fluorescently labeled vesicles. Nuclei and vesicles were segmented independently using IMARIS and their centroid coordinates were exported for spatial analysis.

For each droplet, the geometric center was defined as the origin, and the cartesian coordinates of each nucleus and vesicles were converted into radial distances. The radial position of each detected cell nucleus or vesicles was calculated from its Cartesian coordinates relative to the droplet center:

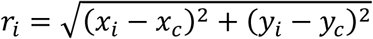

where *x_i_*and *y_i_*are the coordinates of the *i*-th nucleus or vesicle, and *x_c_*and *y_c_*are the coordinates of the droplet centre. The angular coordinate was calculated as:

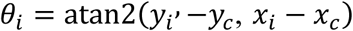

Spatial analyses were performed in RStudio (2026.06.0+242) using custom scripts. Cell and vesicles distributions were visualized using polar scatter plots, and the median radial distance was calculated for each droplet to quantify their spatial localization. Median radial distances from the 10 droplets of each replicate were summarized as whiskers box plots to assess the reproducibility and homogeneity of both cell and vesicles distribution after printing.

### Cell viability

Cell viability was assessed four weeks after encapsulation in RA-GelMA under the culture conditions described above. Samples were stained with Hoechst (**Table 1**) and a LIVE/DEAD assay using calcein-AM and ethidium homodimer-1 (L3224, Thermo Fisher Scientific) according to the manufacturer’s instructions. Hydrogel constructs were sectioned into two halves. For dead controls, samples were incubated in 70% ethanol for 10 min prior to staining. All samples were washed twice in phosphate-buffered saline (PBS) for 10 min and subsequently incubated for 30 min in PBS containing the staining solution. Constructs were then imaged by fluorescence microscopy.

**Table 1.**
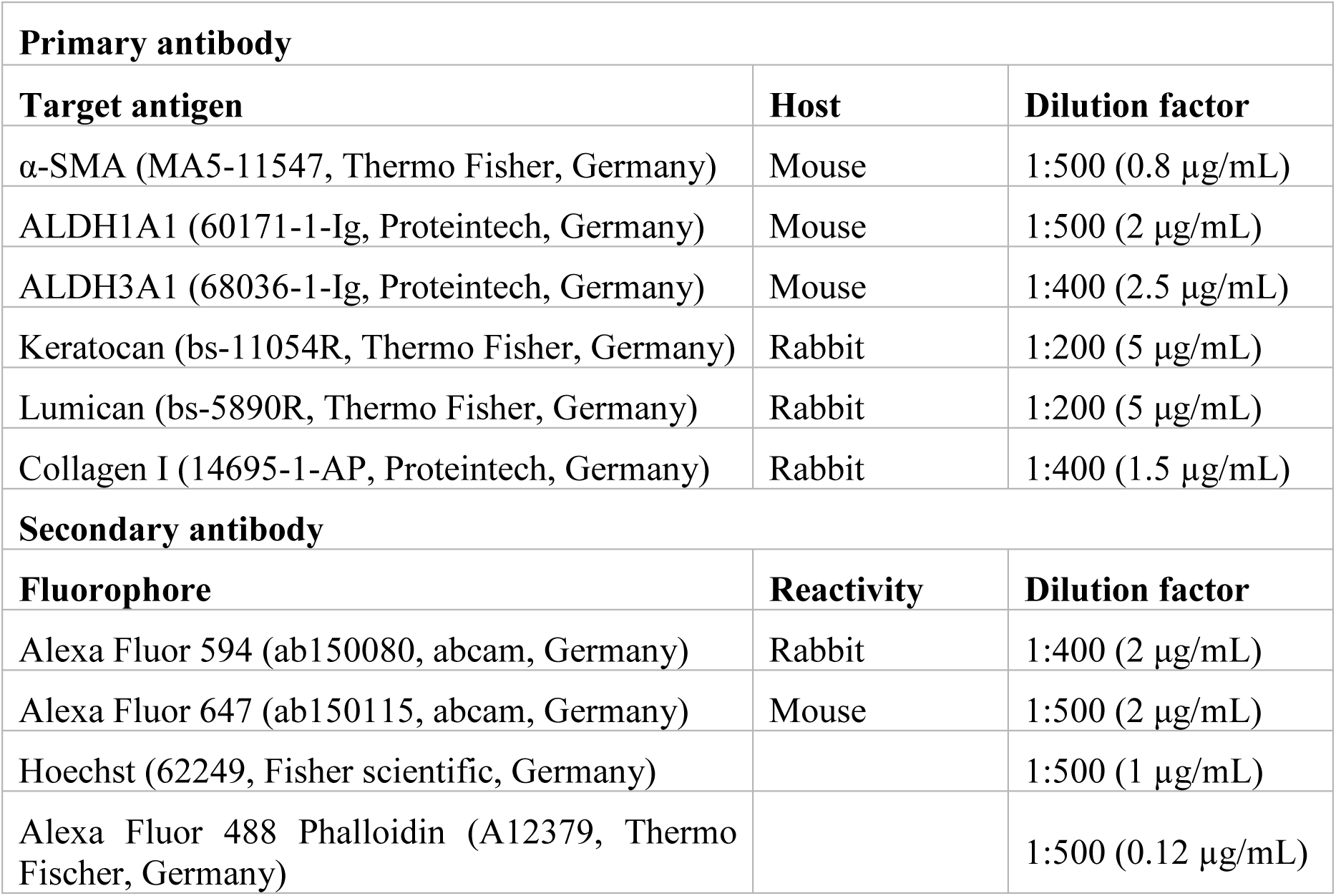
Antibody list and dilution factors.

| **Primary antibody** |  |  |
| --- | --- | --- |
| **Target antigen** | **Host** | **Dilution factor** |
| α-SMA (MA5-11547, Thermo Fisher, Germany) | Mouse | 1:500 (0.8 $\mu\text{g/mL}$) |
| ALDH1A1 (60171-1-Ig, Proteintech, Germany) | Mouse | 1:500 (2 $\mu\text{g/mL}$) |
| ALDH3A1 (68036-1-Ig, Proteintech, Germany) | Mouse | 1:400 (2.5 $\mu\text{g/mL}$) |
| Keratocan (bs-11054R, Thermo Fisher, Germany) | Rabbit | 1:200 (5 $\mu\text{g/mL}$) |
| Lumican (bs-5890R, Thermo Fisher, Germany) | Rabbit | 1:200 (5 $\mu\text{g/mL}$) |
| Collagen I (14695-1-AP, Proteintech, Germany) | Rabbit | 1:400 (1.5 $\mu\text{g/mL}$) |
| **Secondary antibody** |  |  |
| **Fluorophore** | **Reactivity** | **Dilution factor** |
| Alexa Fluor 594 (ab150080, abcam, Germany) | Rabbit | 1:400 (2 $\mu\text{g/mL}$) |
| Alexa Fluor 647 (ab150115, abcam, Germany) | Mouse | 1:500 (2 $\mu\text{g/mL}$) |
| Hoechst (62249, Fisher scientific, Germany) | | 1:500 (1 $\mu\text{g/mL}$) |
| Alexa Fluor 488 Phalloidin (A12379, Thermo Fischer, Germany) | | 1:500 (0.12 $\mu\text{g/mL}$) |

3D image stacks were acquired for each sample (50 slices, step size 2.5 µm). Image analysis was performed using IMARIS. Post-processing included background subtraction and deconvolution. Dead cells were identified as objects exhibiting co-localized Hoechst and ethidium homodimer-1 signals within a defined size range, whereas total cell number was determined based on Hoechst-positive objects. Quantification was performed using spot detection algorithms.

Cell viability was calculated as:

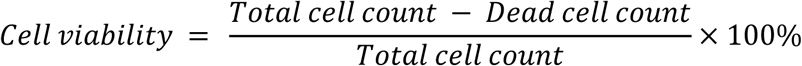

### Rheological characterisation

Samples were allowed to swell to equilibrium in BM-MSC medium at 37 °C for at least 24 h prior to measurement. Rheological characterization was performed using a Discovery Hybrid 20 rheometer (TA Instruments) equipped with an 8 mm parallel plate geometry. Before analysis, hydrogel samples were punched to 8 mm diameter, and excess surface liquid was removed by gentle blotting.

An amplitude sweep (f = 1 Hz, 0.01% ≤ γ ≤ 10%) was conducted to determine the linear viscoelastic region (LVR). The gap height was adjusted individually for each sample to achieve a normal force of 0.1 N. Frequency sweeps were subsequently performed at a constant strain (γ = 1%) over a range of 0.1–100 Hz. All measurements were carried out at 22 °C.

### Quantification of FGF-2 release using ELISA

FGF-2 concentration was quantified by ELISA following the manufacturer’s instructions (900-K08K, PeproTech). Plates were coated overnight with rabbit anti-human FGF capture antibody (1 µg mL⁻¹), blocked with 1% Bovine Serum Albumin (BSA) in PBS, and washed prior to sample addition.

During vesicle preparation, aliquots from the initial FGF-2 solution and wash supernatants were collected. Vesicles were quantified and diluted to 2 × 10⁶ vesicles mL⁻¹. After the third wash, encapsulated FGF-2 was released by treatment with 0.1% Triton X-100 for 10 min at room temperature to induce rupture of the vesicles (n=3, with triplicates tested for each replicate). Samples were diluted to fall within the assay detection range (125–4000 pg mL⁻¹).

A standard serial dilution (125–400 pg mL⁻¹) was performed as described in the manual, standard and samples were loaded in triplicate and incubated for 2 h at room temperature, followed by incubation with biotinylated detection antibody and avidin–HRP. Absorbance was measured at 405 nm and wavelength correction at 650 nm using a microplate reader (FLUOstar Omega, BMG LABTECH). After blank subtraction a standard curve was fitted with a sigmoidal 4PL regression, and the FGF2 concentrations were interpolated and calculated based on the dilution.

### Immunofluorescence staining

Samples were collected and washed three times with PBS supplemented with 1 mM MgCl₂, followed by fixation in 4% paraformaldehyde (47347.9M, VWR) for 20 min at room temperature. After fixation, samples were washed three times with PBS and permeabilized with 0.2% Triton X-100 (1086031000, VWR) for 20 min at room temperature.

Samples were then washed three times with PBS, sectioned into halves, and blocked for 2 h in a blocking solution consisting of 5% donkey serum (P30-0101, Pan-Biotech), 5% goat serum (Pan-Biotech), and 0.1% Triton X-100. Following blocking, samples were incubated with primary antibodies diluted in blocking solution, as detailed in Table 1.

After primary antibody incubation, samples were washed three times with PBS containing 0.2% Tween-20 in the dark. Secondary antibody incubation was then performed for 2 h at room temperature in the dark using fluorophore-conjugated antibodies (goat anti-rabbit Alexa Fluor 594 and goat anti-mouse Alexa Fluor 647). Nuclear and cytoskeletal staining were carried out concurrently using Hoechst and phalloidin-Alexa Fluor 488.

Fluorescence imaging was performed using a confocal microscope (LSM780, Zeiss). For three-dimensional samples, z-stacks were acquired over a depth of 15-20 µm. Image post-processing was conducted using IMARIS, including background subtraction and deconvolution.

### qPCR

RNA isolation from a printed samples was performed following the phenol-chloroform method. Briefly, samples were homogenized with 1 mL TRIzol (15596026, Thermo Fisher, Germany) and snap frozen in liquid nitrogen after digestion. The thawed sample was mixed with 200 μl chloroform, incubated for 2 minutes, and centrifuged for 15 minutes at 12,000 x g at 4 °C. Upon centrifugation, the sample was divided into the phenol-chloroform phase, the interphase, and the upper aqueous phase. The latter that contains the RNA was transferred to a new tube, and it was precipitated by adding 500 μl isopropanol for 10 minutes. The precipitate was then centrifuged for 10 minutes at 12,000 x g. The supernatant was discarded, and the RNA pellet was washed in 1 mL of 75 % ethanol. Then, the sample was centrifuged for 15 minutes at 12,000 x g, and the supernatant was discarded. After airdrying the RNA pellet, it was solubilized in 10 μl RNA-free water.

cDNA synthesis was perform using a synthesis kit (RevertAid First Strand cDNA Synthesis). Gene expression was quantified by qRT-PCR reactions, performed on 1 µg cDNA with the Sso Advanced Universal SYBR Green Supermix (1725270 Bio-Rad, US) and the Bio-Rad CFX Opus 384 Thermocycler.

Fold change was calculated by the 2^−ΔΔCt^ Method.

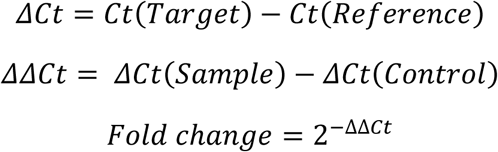

Glyceraldehyde 3-phosphate dehydrogenase (GAPDH) was used as housekeeping gene.

The range of the fold change was then calculated based on the standard deviations (s) from the Ct values.

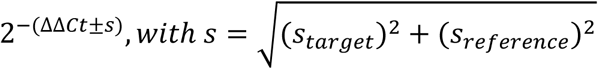

The level of expression of CSK markers before and after differentiation were compared to the level of expression of these markers in BM-MSC cultivated in MSC medium. The level of expression was normalized to BM-MSC. Forward and reverse primers used for qPCR analysis are summarized in Table 2.

**Table 2.**
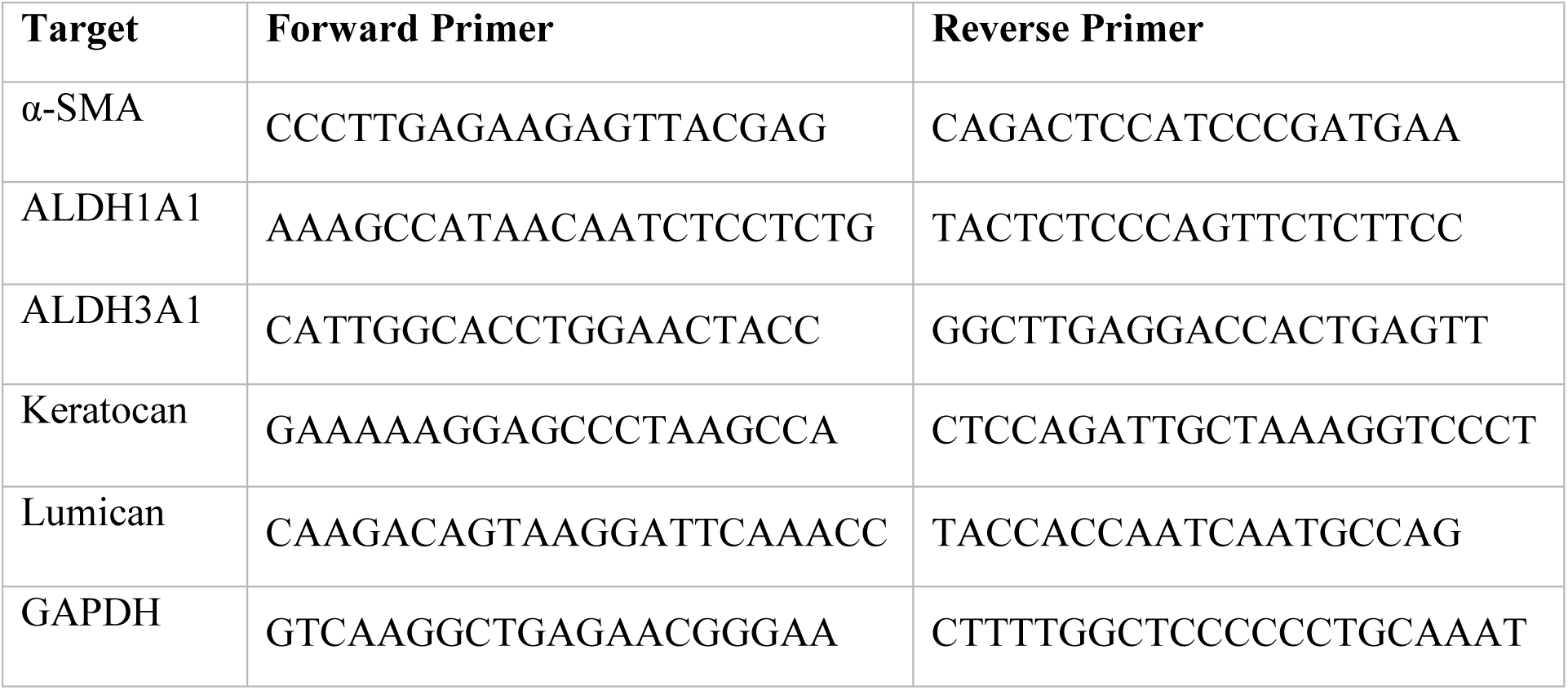
qPCR primers.

### Statistical analysis

All statistical analyses were performed using GraphPad Prism version 10.1.2. Experimental data are presented as mean ± standard deviation (SD), with the corresponding sample size indicated in the respective figure legends. Differences between multiple experimental groups were evaluated using ordinary one-way analysis of variance (ANOVA) followed by Tukey’s multiple-comparison post hoc test. A p-value below 0.05 was considered statistically significant. For comparisons between two matched experimental conditions, statistical significance was assessed using a parametric paired t-test. Levels of statistical significance are indicated as follows: Statistical significance was labeled as \**p* < 0.05, \*\**p* < 0.01, \*\*\**p* < 0.001, and **** p < 0.0001.

## Results

### Growth factor-loaded vesicles remain detectable during long-term 3D culture

FGF-2-loaded and TGF-β3-loaded GUV were incorporated in a 2:1 ratio together with BM-MSC into 30 wt % RA-GelMA constructs (**Figure 1**A). Both fluorescent vesicles populations remained detectable throughout the constructs at days 0, 10, and 17 (**Figure 1**B). The concentration of detectable FGF-2-loaded vesicles decreased from 2.38 ± 0.69 × 10⁶ vesicles mL⁻¹ at day 0 to 1.74 ± 0.38 × 10⁶ vesicles mL⁻¹ at day 10 and 1.52 ± 0.62 × 10⁶ vesicles mL⁻¹ at day 17, although none of the differences between time points reached statistical significance (all adjusted p > 0.05). Similarly, the concentration of TGF-β3-loaded vesicles decreased from 1.24 ± 0.22 × 10⁶ vesicles mL⁻¹ at day 0 to 0.88 ± 0.15 × 10⁶ vesicles mL⁻¹ at day 10 and 0.79 ± 0.33 × 10⁶ vesicles mL⁻¹ at day 17, without significant differences over time (all adjusted p > 0.05). FGF-2-loaded vesicle concentrations were significantly higher than TGF-β3-loaded vesicle concentrations at day 0 (p = 0.0084) and day 10 (p = 0.0362), but not at day 17 (p = 0.0655) (**Figure 1**C). A progressive reduction in the number of detectable vesicles was observed for both populations over time, consistent with gradual loss of vesicle integrity within the hydrogel, interestingly the ratio of 2:1 FGF-2:TGF-β3 vesicles remained constant over the monitored period.

**Figure 1.**
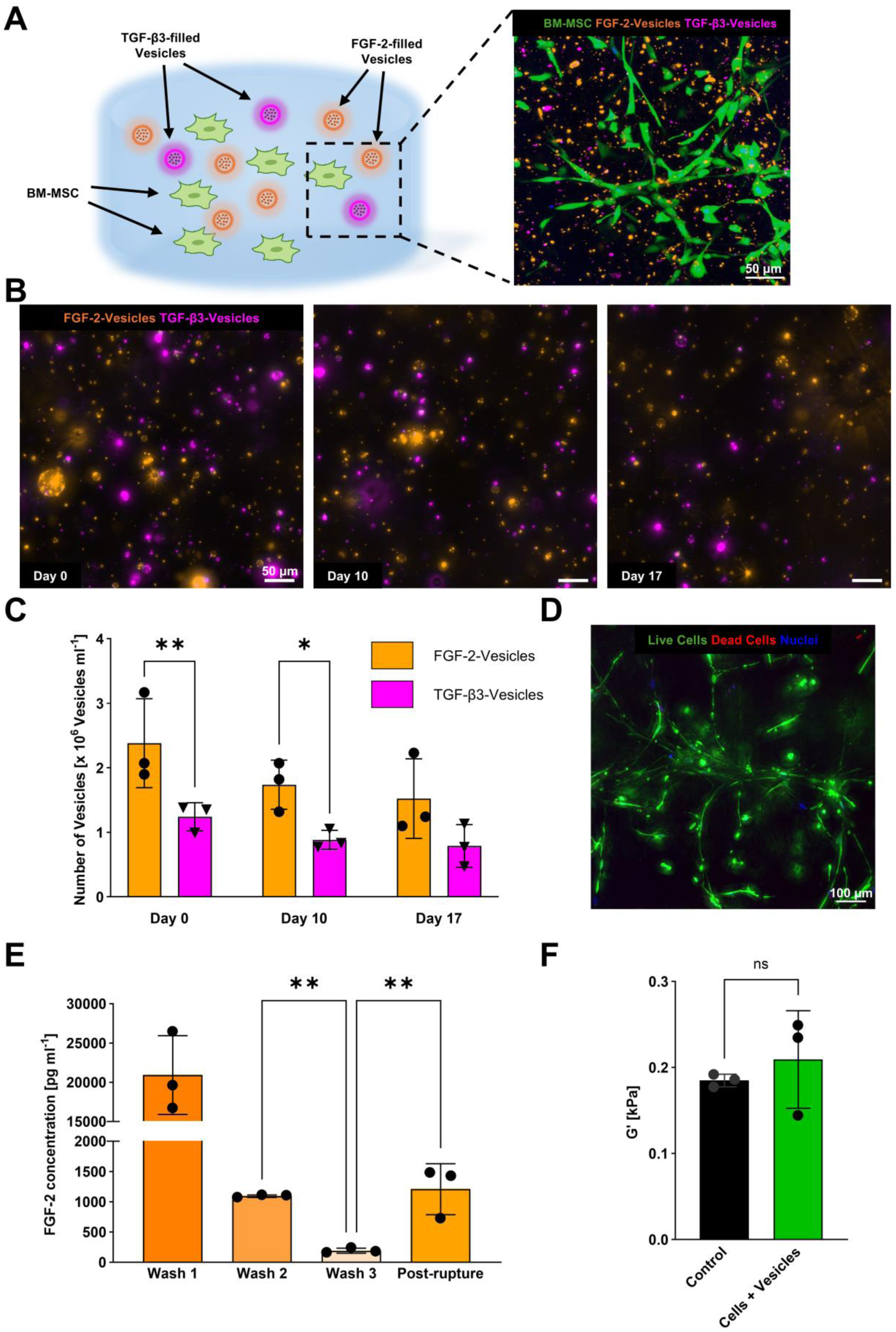
Characterization of growth factor-loaded GUV in BM-MSC-laden RA-GelMA constructs. (A) Schematic representation and representative fluorescence image of BM-MSC (green), FGF-2-loaded GUV (orange), and TGF-β3-loaded GUV (magenta) within RA-GelMA. Scale bar, 50 µm. (B) Representative z-stack maximum intensity projection fluorescence images of FGF-2-loaded GUV and TGF-β3-loaded GUV at days 0, 10, and 17. Scale bars, 50 µm. (C) Quantification of detectable FGF-2-loaded and TGF-β3-loaded GUV during culture. (D) Z-stack projection image of live/dead staining after 28 days, showing live cells in green, dead cells in red, and nuclei in blue. Scale bar, 100 µm. (E) ELISA quantification of FGF-2 in sequential wash fractions and after mediated GUV rupture. (F) Storage modulus (G′) of control RA-GelMA and RA-GelMA containing BM-MSC and GUV. Data are presented as mean ± SD (n = 3). Statistical significance is indicated as *p < 0.05, **p < 0.01, and ns, not significant.

On day 28, live/dead staining showed calcein-positive cells, with limited ethidium homodimer-1 signal (**Figure 1**D). Quantitative image analysis indicated an overall cell viability of 86.5 ± 4.4 %, confirming that BM-MSC remained viable during prolonged culture in RA-GelMA containing growth factor-loaded vesicles.

ELISA measurements showed a progressive decrease in FGF-2 concentration during repeated washing, from 20,918.84 ± 5,011.86 pg mL⁻¹ in wash 1 to 1,092.90 ± 20.08 pg mL⁻¹ in wash 2 and 190.16 ± 41.34 pg mL⁻¹ in wash 3. The FGF-2 concentration in wash 3 was significantly lower than that measured in wash 2 (p = 0.097). Following detergent-induced vesicle lysis, 1,208.09 ± 422.08 pg mL⁻¹ FGF-2 was recovered, which was significantly higher than in wash 3 (p = 0.0054) but not significantly different from wash 2 (p = 0.8377). (**Figure 1**E). These results indicate progressive removal of non-encapsulated FGF-2 during washing, while measurable FGF-2 remained associated with the purified vesicles and could be released following membrane disruption.

Rheological analysis showed comparable storage modulus (G′) values between control RA-GelMA and RA-GelMA containing BM-MSC and vesicles. The storage modulus was 0.185 ± 0.007 kPa for the RA-GelMA control and 0.209 ± 0.057 kPa for RA-GelMA containing BM-MSC and vesicles, with no significant difference between conditions (p = 0.5037). These results indicate that the incorporation of cells and vesicles did not measurably alter the stiffness of RA-GelMA. (**Figure 1**F).

### Drop-on-demand bioprinting preserves the spatial distribution of BM-MSC and GUV

Printability of the RA-GelMA hydrogel was assessed for several polymer concentrations, with the 30 wt % formulation producing reproducible droplets in 7 of the 16 tested pressure and valve-opening combinations. This concentration was selected because 30 wt % RA-GelMA had previously been established for corneal biofabrication[32, 33, 54], although, to our knowledge, it had not yet been processed by drop-on-demand bioprinting. Reliable printing required higher pressures and longer valve-opening times, while droplet mass remained tunable within the printable range and increased with valve-opening time, ranging from 0.17 ± 0.01 mg to 0.69 ± 0.11 mg per droplet. These results identified a suitable processing window for the bioprinting of 30 wt % RA-GelMA while preserving control over droplet deposition, with the complete printability matrix and corresponding droplet mass values provided in the Supplementary Information (**Figure S1, Table S1**).

Drop-on-demand bioprinting generated individual RA-GelMA droplets in which both BM-MSC and fluorescent vesicles were readily detected (**Figure 2**A). Cell nuclei and vesicles were segmented independently, and the resulting masks were used to extract centroid coordinates within the defined droplet boundary (**Figure 2**B). Polar visualization showed that both populations occupied central and peripheral regions of the printed droplets without a dominant angular accumulation (**Figure 2**C).

**Figure 2.**
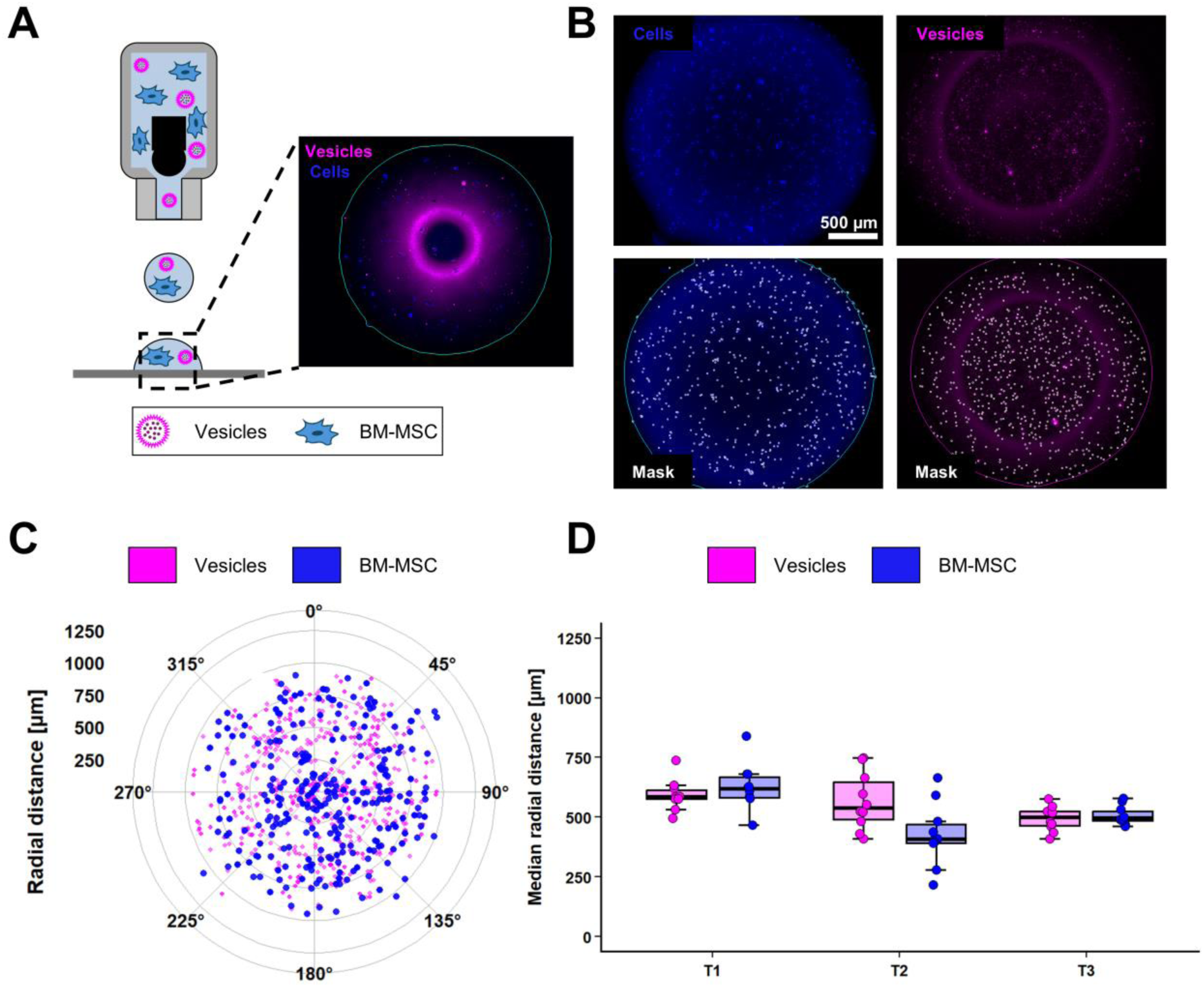
Spatial distribution of BM-MSC and GUV following drop-on-demand bioprinting. (A) Schematic representation of droplet printing and a representative fluorescence image of a printed RA-GelMA droplet containing 1 ×10⁶ BM-MSC mL⁻¹ (blue) and 1 ×10⁶ vesicles mL⁻¹ (magenta). (B) Representative fluorescence images and corresponding segmentation masks used to identify cell nuclei and GUV within the droplet boundary in separate bioinks. Scale bar, 500 µm. (C) Representative polar scatter plot showing the angular and radial positions of BM-MSC and GUV relative to the droplet center. (D) Median radial distance of BM-MSC and GUV in 10 individual droplets from each independent experiment (T1-T3). Individual points represent droplets; boxes indicate the median and interquartile range.

The spatial distribution of cells and vesicles was assessed from 30 droplets, corresponding to three independent replicates with 10 droplets per replicate. A total of 16,474 cells and 16,505 vesicles were detected across 30 droplets. This corresponded to an average of 549.1 ± 130.5 cells and 550.2 ± 240.4 vesicles per droplet (**Figure 2**B). Consistent overlap between the two object populations was observed, indicating that vesicles and cells were dispersed throughout similar regions of the printed droplets (**Figure 2**C).

The radial distribution nevertheless varied between replicates (**Figure 2**D). In T1, the mean droplet-level median radial distance was 593.0 ± 64.4 µm for vesicles and 627.8 ± 95.8 µm for cells, indicating comparable distributions. In T2, the mean droplet-level median radial distance was 567.6 ± 119.8 µm for vesicles, compared with 427.9 ± 131.5 µm for cells. In T3, the distributions again overlapped, with median radial distances of 494.8 ± 51.1 µm for vesicles and 507.4 ± 39.1 µm for cells.

Across all 30 droplets, the mean droplet-level median radial distance was 551.8 ± 91.3 µm for vesicles and 521.0 ± 125.2 µm for cells. Overall, the polar distribution analysis demonstrated that the vesicles remained spatially associated with the cell-containing regions following printing, although replicate-dependent variations in their relative radial distributions were observed.

### GUV-mediated growth factor delivery supports keratocyte-associated protein expression after bioprinting

Following a 3-day spreading phase, BM-MSC were differentiated for 14 days using either soluble growth factor supplementation or vesicle-mediated delivery of FGF-2 and TGF-β3 in growth factor-free differentiation medium (**Figure 3**A). Representative immunofluorescence images showed clear differences between undifferentiated BM-MSC and the differentiated conditions (**Figure 3**B). BM-MSC cultured under basal conditions showed no detectable ALDH1A1, ALDH3A1, Lumican, or Keratocan signal, with only weak intracellular Collagen I staining and no detectable α-SMA. In contrast, conventionally differentiated CSK-MSC and vesicle-differentiated-MSC (GUV-MSC) showed positive staining for ALDH1A1, ALDH3A1, Lumican, Keratocan, and Collagen I. The same keratocyte-associated markers remained detectable in GUV-MSC following drop-on-demand bioprinting, while α-SMA remained undetectable or low across all conditions. Single signals for all immunofluorescence stainings are provided in supplementary section (**Figure S2** to **Figure S5**) Printed BM-MSC and CSK-MSC are also provided in supplementary section (**Figure S6** and **Figure S7**)

**Figure 3.**
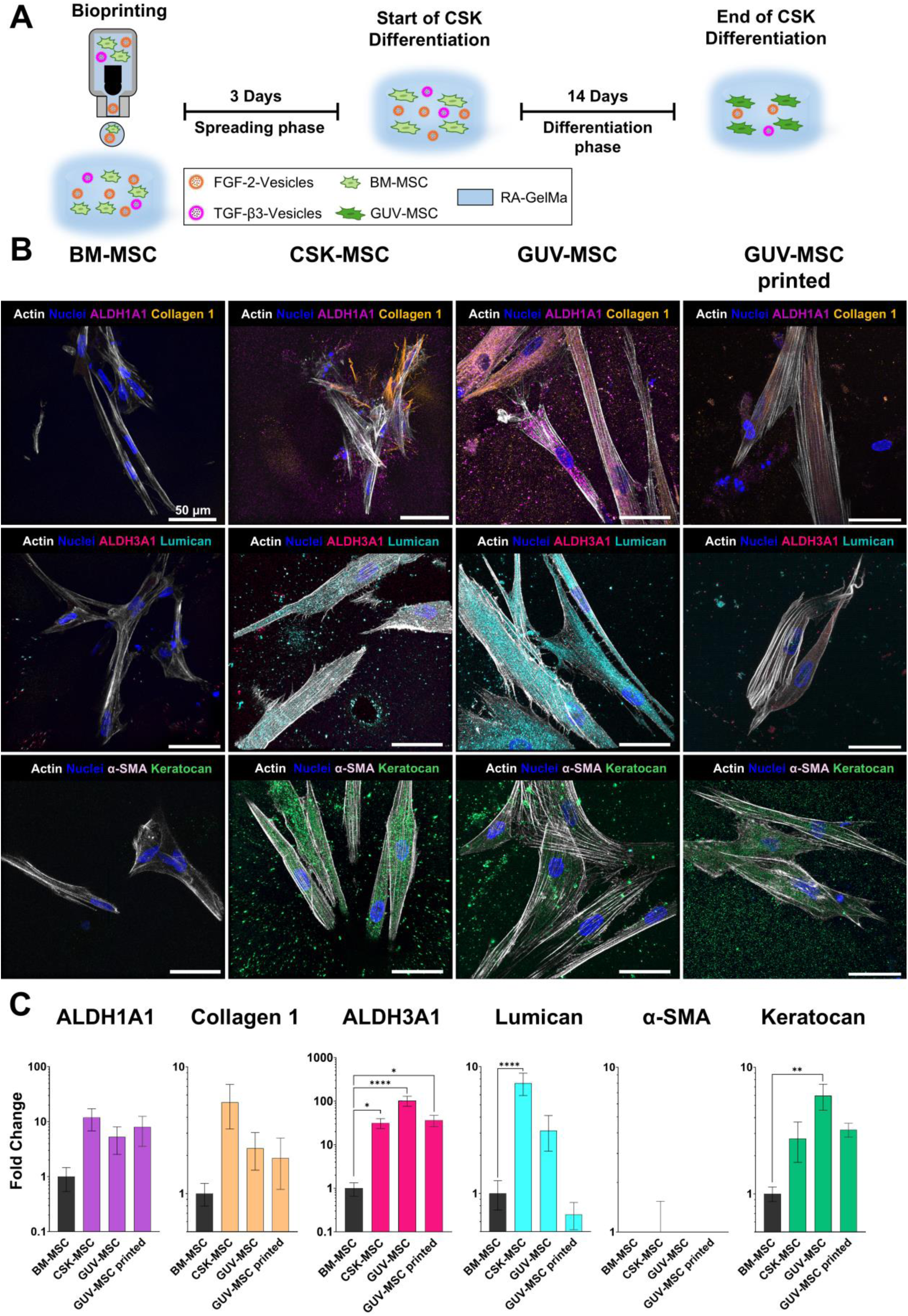
GUV-mediated growth factor delivery induces keratocyte-associated marker expression and remains effective following drop-on-demand bioprinting. (A) Experimental workflow for bioprinting, the 3-day spreading phase, and the 14-day keratocyte differentiation phase. (B) Representative immunofluorescence images of undifferentiated BM-MSC, conventionally differentiated CSK-MSC, GUV-MSC differentiated using FGF-2-loaded and TGF-β3-loaded GUV, and printed GUV-MSC. Constructs were stained for ALDH1A1 (magenta), Collagen I (orange), ALDH3A1 (pink), Lumican (cyan), α-SMA (lilac), and Keratocan (green). Actin filaments were stained with phalloidin (white) and nuclei with Hoechst (blue). Scale bars, 50 µm. (C) Quantification of immunofluorescence signal intensity, expressed as fold change relative to undifferentiated BM-MSC. Data are presented as mean ± SD (n = 3 technical replicates from a pooled sample of three donors). Statistical significance is indicated as *p < 0.05, **p < 0.01, ***p < 0.001, and ****p < 0.0001.

Quantification of the immunofluorescence signal is presented in **Figure 3**C. ALDH1A1 expression increased to 11.99 ± 5.19-fold in CSK-MSC, 5.28 ± 2.74-fold in GUV-MSC, and 8.04 ± 4.49-fold in printed GUV-MSC relative to undifferentiated BM-MSC. Collagen I expression increased to 5.25 ± 2.02-fold, 2.28 ± 0.75-fold, and 1.91 ± 0.83-fold, respectively.

ALDH3A1 expression was significantly increased relative to undifferentiated BM-MSC in CSK-MSC (31.65 ± 8.23-fold, p = 0.0402), GUV-MSC (103.15 ± 26.86-fold, p < 0.0001), and printed GUV-MSC (36.64 ± 10.73-fold, p = 0.0190). ALDH3A1 induction was also higher in GUV-MSC than in CSK-MSC (p = 0.0002) and printed GUV-MSC (p = 0.0004).

Lumican expression increased significantly in CSK-MSC (7.41 ± 1.48-fold, p < 0.0001), whereas the increases measured in GUV-MSC (3.14 ± 0.98-fold) and printed GUV-MSC (0.68 ± 0.17-fold) did not reach significance relative to undifferentiated BM-MSC. Keratocan expression increased to 2.73 ± 0.96-fold in CSK-MSC, 5.96 ± 1.39-fold in GUV-MSC, and 3.21 ± 0.40-fold in printed GUV-MSC, with a significant increase observed in the GUV-MSC condition (p = 0.0100). α-SMA remained low in CSK-MSC (0.87 ± 0.66-fold), GUV-MSC (0.38 ± 0.15-fold), and printed GUV-MSC (0.51 ± 0.26-fold), with no significant differences relative to undifferentiated BM-MSC.

### GUV-mediated differentiation promotes a keratocyte-like transcriptional profile after bioprinting

The expression of keratocyte-associated genes was quantified by qRT-PCR following differentiation in bioprinted RA-GelMA constructs (**Figure 4**).

**Figure 4.**
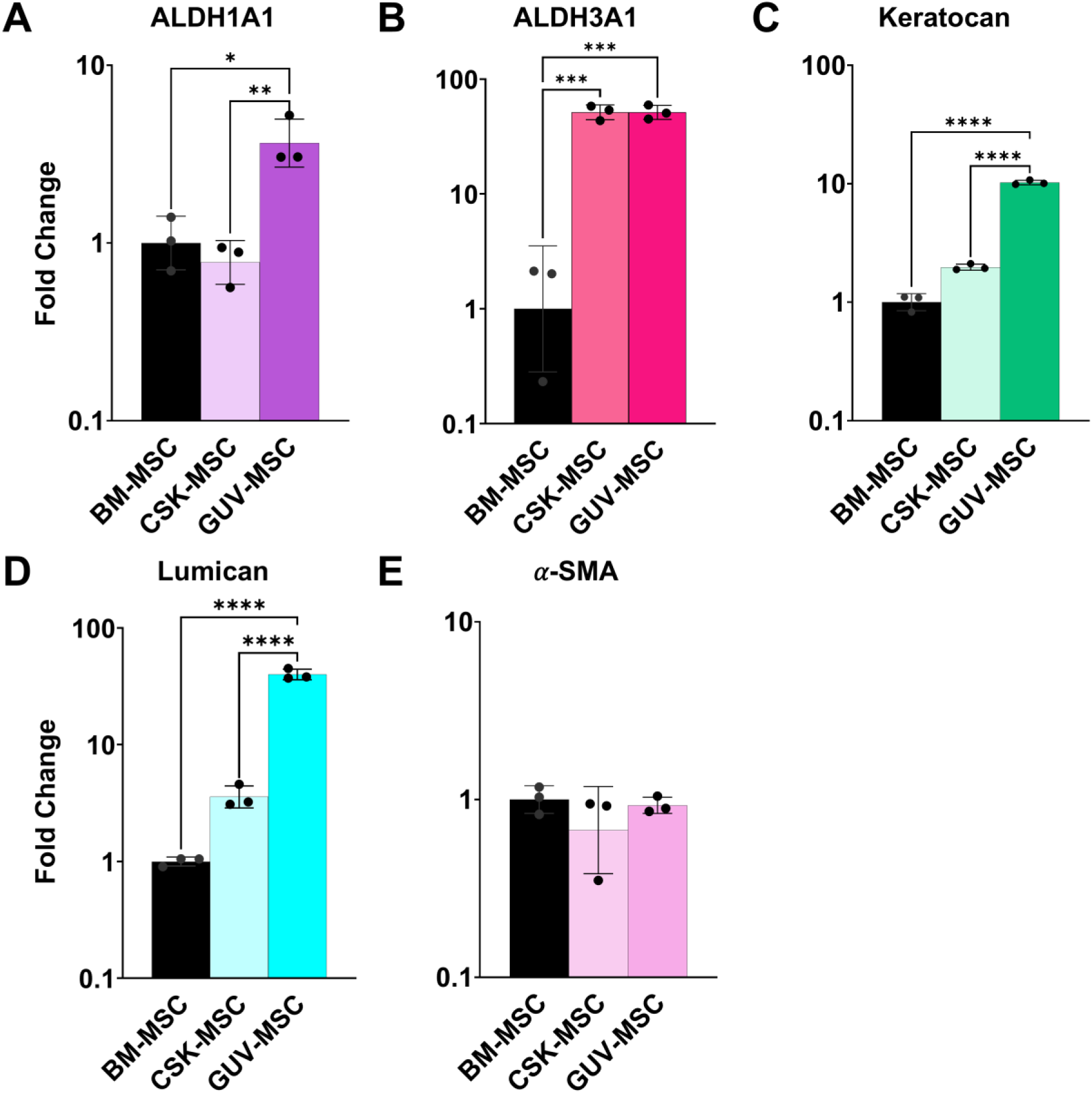
GUV-mediated differentiation promotes keratocyte-associated gene expression without α-SMA induction after bioprinting. Relative gene expression of (A) ALDH1A1, (B) ALDH3A1, (C) Keratocan, (D) Lumican, and (E) α-SMA in undifferentiated BM-MSC, conventionally differentiated CSK-MSC, and GUV-MSC following drop-on-demand bioprinting in RA-GelMA. Gene expression was normalized to GAPDH and expressed as fold change relative to BM-MSC. Data are presented as mean ± SD from three biological replicates. Statistical significance is indicated as *p < 0.05, **p < 0.01, ***p < 0.001, ****p < 0.0001, and ns, not significant.

ALDH1A1 expression was 1.04 ± 0.35-fold in BM-MSC and 0.80 ± 0.21-fold in CSK-MSC. GUV-MSC showed an ALDH1A1 expression of 3.77 ± 1.26-fold, which was significantly higher than both BM-MSC (p = 0.0110) and CSK-MSC (p = 0.0074).

ALDH3A1 expression was 1.46 ± 1.06-fold in BM-MSC, 51.82 ± 7.46-fold in CSK-MSC, and 51.74 ± 7.34-fold in GUV-MSC. Expression was significantly higher in both CSK-MSC and GUV-MSC than in BM-MSC (p = 0.0001 for both comparisons), while no significant difference was detected between the two differentiated conditions.

Keratocan expression was 1.01 ± 0.16-fold in BM-MSC and increased to 1.98 ± 0.12-fold in CSK-MSC (p = 0.0149). A greater increase was detected in GUV-MSC, which reached 10.25 ± 0.46-fold and was significantly higher than both BM-MSC and CSK-MSC (p < 0.0001 for both comparisons).

Lumican expression was 1.00 ± 0.09-fold in BM-MSC and 3.63 ± 0.83-fold in CSK-MSC, with no significant difference between these conditions. GUV-MSC reached 40.11 ± 4.29-fold, which was significantly higher than both BM-MSC and CSK-MSC (p < 0.0001 for both comparisons). α-SMA expression remained low across all conditions, with values of 1.01 ± 0.18-fold in BM-MSC, 0.74 ± 0.34-fold in CSK-MSC, and 0.93 ± 0.10-fold in GUV-MSC, and no significant differences were detected.

Overall, GUV-MSC reached ALDH3A1 expression levels comparable to conventionally differentiated CSK-MSC, while ALDH1A1, Lumican, and Keratocan expression were higher in GUV-MSC. The absence of α-SMA induction further supported acquisition of a keratocyte-like transcriptional profile without myofibroblastic activation.

## Discussion

The present study investigated whether growth factor-loaded vesicles could be integrated into a bioink to provide locally available biochemical cues for BM-MSC differentiation following drop-on-demand bioprinting. The results support the feasibility of combining BM-MSC, FGF-2- and TGF-β3-loaded vesicles within a single printable formulation. Following purification, FGF-2 remained detectable after vesicle rupture, they were progressively reduced in number during culture, and BM-MSC remained viable within the resulting constructs. Moreover, keratocyte-associated protein and gene expressions were detected following vesicle-mediated differentiation, including bioprinted samples. These findings establish synthetic cell-like vesicles as compatible components of a 3D biofabrication system and indicate that vesicle-mediated delivery can provide sufficient biochemical stimulation to promote a keratocyte-like phenotype without repeated supplementation of soluble growth factors.

The biochemical characterization of the vesicle preparation provided evidence that non-encapsulated FGF-2 was efficiently removed during the purification step. The reduction in FGF-2 concentration between the first and second washes, followed by concentrations close to the detection limit in the third wash, indicated that freely dissolved growth factor was largely eliminated. Even further reduced when incorporating the vesicles into a bioink. The recovery of FGF-2 after detergent-mediated vesicle rupture further confirmed that the final preparation retained measurable growth factor. Although FGF-2 detectability after vesicle rupture does not by itself demonstrate that the released protein retained its full biological activity, the following differentiation response provides indirect functional evidence that the vesicle-containing formulation generated biologically relevant conditions.

A progressive decrease in the number of detectable FGF-2- and TGF-β3-loaded GUV was observed between day 0 and day 17. Gradual vesicle destabilization or rupture within hydrogels could support time-dependent cargo availability during the differentiation period[22, 55, 56]. The successful printing of 30 wt % RA-GelMA bioink containing BM-MSC and growth factor-loaded vesicles demonstrated the compatibility of the complete formulation with drop-on-demand bioprinting. Both cells and vesicles were consistently detected throughout individual droplets, indicating effective co-deposition. This spatial proximity supports the possibility that progressive vesicle rupture may maintain growth factor availability to the surrounding cells during differentiation period.

From a biological perspective, BM-MSC maintained high viability over 28 days in 3D RA-GelMA, confirming the cytocompatibility of both the hydrogel and vesicle incorporation. Following 14 days of differentiation, immunofluorescence analysis demonstrated expression of keratocyte-associated markers, including ALDH1A1, ALDH3A1, lumican, and keratocan, together with collagen I deposition and absence of α-SMA expression. These findings are indicative of differentiation towards a keratocyte-like phenotype without transition to a fibroblastic state, which is critical for preserving corneal transparency.

Following drop-on-demand bioprinting, qRT-PCR analysis showed that GUV-MSC maintained a keratocyte-associated transcriptional profile. ALDH3A1 expression was comparable to conventionally differentiated CSK-MSC, whereas ALDH1A1, lumican, and keratocan expression were significantly higher in GUV-MSC than in both BM-MSC and CSK-MSC. α-SMA expression remained low and unchanged between conditions. As differentiation was achieved in growth factor-free medium, these results indicate that the incorporated GUV provided sufficient biochemical cues to support keratocyte-like differentiation after printing without repeated supplementation with soluble growth factors.

Previous corneal bioprinting studies have primarily focused on the direct encapsulation of primary keratocytes within collagen- or GelMA-based bioinks, demonstrating high viability, stromal marker maintenance, transparency, and mimicking the native corneal shape[57–59]. More recent approaches have incorporated limbus-derived MSC into cornea-derived extracellular matrix bioinks[60] or used biomaterial composition and scaffold architecture to promote keratocyte phenotype maintenance[61]. However, primary keratocytes remain difficult to expand without phenotypic loss[62, 63], whereas BM-MSC are easily collectable and their expansion *in vitro* is widely protocolized[64–66].

The present system addresses both limitations by combining an expandable BM-MSC source with growth factor-loaded vesicles directly embedded within a printable RA-GelMA bioink. In contrast to approaches that print an already differentiated cell population or rely on repeated soluble-factor supplementation, the biochemical cues required for differentiation are co-deposited with the cells and remain locally available within each printed droplet. Differentiation in growth factor-free medium after printing therefore represents a key advantage of this system for settings in which repeated factor administration would be impractical, particularly *in situ* biofabrication. Rather than being superior to existing corneal bioinks, this strategy provides a functional advantage by integrating cell delivery, localized biochemical instruction, and post-printing differentiation within a single formulation.

Several additional limitations should be considered. FGF-2 was quantified following purification and vesicle rupture, whereas TGF-β3 encapsulation was not independently quantified. Vesicle counting was restricted to vesicles larger than 8 µm, as smaller vesicles were expected to contain less cargo; consequently, their contribution to the total vesicle population was not counted. The lamellarity, cargo loading variability, and stability of encapsulated growth factor in the vesicles during the differentiation period were not characterized. Donor-derived BM-MSC were pooled, which reduced inter-donor variability within the experiments but prevented evaluation of donor-specific responses.

Future studies should therefore quantify the release and biological activity of both FGF-2 and TGF-β3 throughout the differentiation time, determine vesicle loading efficiency and size-dependent stability, and establish the relationship between local vesicle density and cell response. Independent donor experiments should be included to assess reproducibility across BM-MSC populations. The resulting tissue should also be evaluated for collagen organization, proteoglycan deposition, transparency, refractive properties, and integration within corneal stromal models.

## Conclusion

Engineered cell-like vesicles embedded within RA-GelMA hydrogels enable sustained and localized growth factor delivery, supporting keratocyte differentiation of BM-MSC in both cast and bioprinted 3D constructs. This approach overcomes the need for repeated supplementation and remains compatible with drop-on-demand bioprinting, highlighting its potential for *in situ* corneal bioprinting. More broadly, programmable vesicle-based microenvironments offer a platform to precisely regulate cell behavior within engineered tissues, with the potential to be adapted across a wide range of tissue-specific applications.

## AI use statement

During the preparation of this work, ChatGPT (OpenAI) and Claude (Anthropic) were used to assist in generating and refining R scripts for data processing, polar coordinate transformation, statistical data organization, and graphical visualization. All generated code was reviewed, adapted, and validated by the authors, and the resulting outputs were checked against the original datasets. The tools were not used to independently interpret the results or formulate scientific conclusions. The authors assume full responsibility for the accuracy and integrity of the analyses presented.

## Acknowledgements

This work was supported by the *Bundesministerium für Forschung, Technologie und Raumfahrt* (BMFTR, Grant Number 13XP5135 to DFDC). This work was supported by the Deutsche Forschungsgemeinschaft (DFG, German Research Foundation) under Germany’s Excellence Strategy via the Excellence Cluster 3D Matter Made to Order (EXC-2082/1–390761711 to DFDC). The authors thank Prof. Dr. Henrik Kaessmann and his group for the use of their qPCR machine.

## Data Availability Statement

The data that support the findings of this study are available from the corresponding author upon reasonable request.

## Author contributions

A.T. and O.T. designed the study, performed the experiments, analyzed the data, and wrote the original manuscript, they contributed equally to this work. A.J.A. did vesicle production, 3D culture, IF staining. P.S. did GelMA production, bioprinting experiments. C.R.W. did ELISA and 3D cultures. A.F. did ELISA and design of the study. M.W. did 2D cultures. F.D. did GelMA production, rheology, and revised the manuscript. D.F.D.C. supervised the study, acquired funding, and contributed to writing and editing. All authors reviewed and approved the final manuscript.

## References

[1] M. Abdul-Al, G. K. Kyeremeh, M. Saeinasab, S. Heidari Keshel, and F. Sefat, “Stem Cell Niche Microenvironment: Review,” (in eng), Bioengineering (Basel), vol. 8, no. 8, Jul 28 2021, doi: 10.3390/bioengineering8080108.

[2] B. Choi, D. Kim, I. Han, and S. H. Lee, “Microenvironmental Regulation of Stem Cell Behavior Through Biochemical and Biophysical Stimulation,” (in eng), Adv Exp Med Biol, vol. 1064, pp. 147–160, 2018, doi: 10.1007/978-981-13-0445-3_9.

[3] F. Gattazzo, A. Urciuolo, and P. Bonaldo, “Extracellular matrix: A dynamic microenvironment for stem cell niche,” Biochimica et Biophysica Acta (BBA) - General Subjects, vol. 1840, no. 8, pp. 2506–2519, 2014/08/01/ 2014, doi: 10.1016/j.bbagen.2014.01.010.

[4] M. A. Kinney and T. C. McDevitt, “Emerging strategies for spatiotemporal control of stem cell fate and morphogenesis,” (in eng), Trends Biotechnol, vol. 31, no. 2, pp. 78–84, Feb 2013, doi: 10.1016/j.tibtech.2012.11.001.

[5] R. Subbiah and R. E. Guldberg, “Materials Science and Design Principles of Growth Factor Delivery Systems in Tissue Engineering and Regenerative Medicine,” Advanced Healthcare Materials, vol. 8, no. 1, p. 1801000, 2019, doi: 10.1002/adhm.201801000.

[6] J. Temple, E. Velliou, M. Shehata, R. Lévy, and P. Gupta, “Current strategies with implementation of three-dimensional cell culture: the challenge of quantification,” Interface Focus, vol. 12, no. 5, 2022, doi: 10.1098/rsfs.2022.0019.

[7] A. M. Leferink, D. Santos, M. Karperien, R. K. Truckenmüller, C. A. van Blitterswijk, and L. Moroni, “Differentiation capacity and maintenance of differentiated phenotypes of human mesenchymal stromal cells cultured on two distinct types of 3D polymeric scaffolds,” Integrative Biology, vol. 7, no. 12, pp. 1574–1586, 2015, doi: 10.1039/c5ib00177c.

[8] S. Raghavan, C. J. Shen, R. A. Desai, N. J. Sniadecki, C. M. Nelson, and C. S. Chen, “Decoupling diffusional from dimensional control of signaling in 3D culture reveals a role for myosin in tubulogenesis,” (in eng), J Cell Sci, vol. 123, no. Pt 17, pp. 2877–83, Sep 1 2010, doi: 10.1242/jcs.055079.

[9] O. Habanjar, M. Diab-Assaf, F. Caldefie-Chezet, and L. Delort, “3D Cell Culture Systems: Tumor Application, Advantages, and Disadvantages,” (in eng), Int J Mol Sci, vol. 22, no. 22, Nov 11 2021, doi: 10.3390/ijms222212200.

[10] B. M. Baker and C. S. Chen, “Deconstructing the third dimension – how 3D culture microenvironments alter cellular cues,” Journal of Cell Science, vol. 125, no. 13, pp. 3015–3024, 2012, doi: 10.1242/jcs.079509.

[11] W. Chen, P. Wu, C. Jin, Y. Chen, C. Li, and H. Qian, “Advances in the application of extracellular vesicles derived from three-dimensional culture of stem cells,” (in eng), J Nanobiotechnology, vol. 22, no. 1, p. 215, May 1 2024, doi: 10.1186/s12951-024-02455-y.

[12] X. Yuan et al., “Engineering extracellular vesicles by three-dimensional dynamic culture of human mesenchymal stem cells,” (in eng), J Extracell Vesicles, vol. 11, no. 6, p. e12235, Jun 2022, doi: 10.1002/jev2.12235.

[13] M. Casajuana Ester and R. M. Day, “Production and Utility of Extracellular Vesicles with 3D Culture Methods,” (in eng), Pharmaceutics, vol. 15, no. 2, Feb 16 2023, doi: 10.3390/pharmaceutics15020663.

[14] A. Poerio, J. o. F. Mano, and F. Cleymand, “Advanced 3D Printing Strategies for the Controlled Delivery of Growth Factors,” ACS Biomaterials Science & Engineering, vol. 9, no. 12, pp. 6531–6547, 2023, doi: 10.1021/acsbiomaterials.3c00873.

[15] L. M. Caballero Aguilar, S. M. Silva, and S. E. Moulton, “Growth factor delivery: Defining the next generation platforms for tissue engineering,” Journal of Controlled Release, vol. 306, pp. 40–58, 2019/07/28/ 2019, doi: 10.1016/j.jconrel.2019.05.028.

[16] T. T. Goodman, C. P. Ng, and S. H. Pun, “3-D tissue culture systems for the evaluation and optimization of nanoparticle-based drug carriers,” (in eng), Bioconjug Chem, vol. 19, no. 10, pp. 1951–9, Oct 2008, doi: 10.1021/bc800233a.

[17] D. A. Shah, S. J. Kwon, S. S. Bale, A. Banerjee, J. S. Dordick, and R. S. Kane, “Regulation of stem cell signaling by nanoparticle-mediated intracellular protein delivery,” (in eng), Biomaterials, vol. 32, no. 12, pp. 3210–9, Apr 2011, doi: 10.1016/j.biomaterials.2010.11.077.

[18] S. Bruno, G. Chiabotto, E. Favaro, M. C. Deregibus, and G. Camussi, “Role of extracellular vesicles in stem cell biology,” American Journal of Physiology-Cell Physiology, vol. 317, no. 2, pp. C303-C313, 2019, doi: 10.1152/ajpcell.00129.2019.

[19] M. Kou et al., “Mesenchymal stem cell-derived extracellular vesicles for immunomodulation and regeneration: a next generation therapeutic tool?,” Cell Death & Disease, vol. 13, no. 7, p. 580, 2022/07/04 2022, doi: 10.1038/s41419-022-05034-x.

[20] W. Cheng et al., “Engineered Extracellular Vesicles: A potential treatment for regeneration,” iScience, vol. 26, no. 11, 2023, doi: 10.1016/j.isci.2023.108282.

[21] Z. Pan et al., “Extracellular Vesicles in Tissue Engineering: Biology and Engineered Strategy,” Advanced Healthcare Materials, vol. 11, no. 21, p. 2201384, 2022, doi: 10.1002/adhm.202201384.

[22] O. Thaden et al., “Bioprinting of Synthetic Cell-like Lipid Vesicles to Augment the Functionality of Tissues after Manufacturing,” ACS Synthetic Biology, vol. 13, no. 8, pp. 2436–2446, 2024/8// 2024, doi: 10.1021/ACSSYNBIO.4C00137/ASSET/IMAGES/LARGE/SB4C00137_0006.JPEG.

[23] R. Cochereau, V. Maffeis, E. C. dos Santos, E. Lörtscher, and C. G. Palivan, “Polymeric Giant Unilamellar Vesicles with Integrated DNA-Origami Nanopores: An Efficient Platform for Tuning Bioreaction Dynamics Through Controlled Molecular Diffusion,” Advanced Functional Materials, vol. 33, no. 48, p. 2304782, 2023, doi: 10.1002/adfm.202304782.

[24] A. Chen, S. Gat, L. Ohana, E. Yekymov, Y. Tsori, and A. Bernheim-Groswasser, “Strategy for Generating Giant Unilamellar Vesicles with Tunable Size Using the Modified cDICE Method,” (in eng), ACS Synth Biol, vol. 14, no. 7, pp. 2597–2608, Jul 18 2025, doi: 10.1021/acssynbio.5c00026.

[25] Y. Shao, J. Sang, and J. Fu, “On human pluripotent stem cell control: The rise of 3D bioengineering and mechanobiology,” (in eng), Biomaterials, vol. 52, pp. 26–43, Jun 2015, doi: 10.1016/j.biomaterials.2015.01.078.

[26] M. F. Simsek and E. M. Özbudak, “Patterning principles of morphogen gradients,” Open Biology, vol. 12, no. 10, 2022, doi: 10.1098/rsob.220224.

[27] B. J. O’Grady, D. A. Balikov, E. S. Lippmann, and L. M. Bellan, “Spatiotemporal Control of Morphogen Delivery to Pattern Stem Cell Differentiation in Three-Dimensional Hydrogels,” (in eng), Curr Protoc Stem Cell Biol, vol. 51, no. 1, p. e97, Dec 2019, doi: 10.1002/cpsc.97.

[28] L. P. Guérin et al., “The Human Tissue-Engineered Cornea (hTEC): Recent Progress,” (in eng), Int J Mol Sci, vol. 22, no. 3, Jan 28 2021, doi: 10.3390/ijms22031291.

[29] A. Zhang, W. Zhang, L. J. Backman, and J. Chen, “Advances in Regulatory Strategies of Differentiating Stem Cells towards Keratocytes,” (in eng), Stem Cells Int, vol. 2022, p. 5403995, 2022, doi: 10.1155/2022/5403995.

[30] J. Wu, Y. Du, M. M. Mann, E. Yang, J. L. Funderburgh, and W. R. Wagner, “Bioengineering organized, multilamellar human corneal stromal tissue by growth factor supplementation on highly aligned synthetic substrates,” (in eng), Tissue Eng Part A, vol. 19, no. 17-18, pp. 2063–75, Sep 2013, doi: 10.1089/ten.TEA.2012.0545.

[31] L. E. Sidney and A. Hopkinson, “Corneal keratocyte transition to mesenchymal stem cell phenotype and reversal using serum-free medium supplemented with fibroblast growth factor-2, transforming growth factor-â3 and retinoic acid,” (in eng), J Tissue Eng Regen Med, vol. 12, no. 1, pp. e203-e215, Jan 2018, doi: 10.1002/term.2316.

[32] A. Taoum et al., “3D Differentiation of Bone-Marrow Derived Mesenchymal Stromal Cells into the Keratocyte Lineage for Corneal Bioprinting,” Advanced Healthcare Materials, vol. 14, no. 28, p. 2405073, 2025, doi: 10.1002/adhm.202405073.

[33] A. Taoum et al., “Toward clinically relevant automated corneal biomanufacturing with human-derived FBS alternatives,” Scientific Reports, vol. 16, no. 1, p. 18874, 2026/06/17 2026, doi: 10.1038/s41598-026-58401-5.

[34] Y. Jian, F. Dehli, M. Wisbar, A. Taoum, and D. Duarte Campos, “In situ bioprinting: bioprinting methods, bioinks, cell sources & advanced bioprinting strategies,” Biofabrication, vol. 18, no. 1, p. 012007, 2026/02/09 2026, doi: 10.1088/1758-5090/ae3cc3.

[35] S. Abraham, S. P. Rangaswamy, and A. Chinnaiah, “Evaluation of recombinant human vascular endothelial growth factor VEGF121-loaded poly-l-lactide microparticles as a controlled release delivery system,” (in eng), Turk J Biol, vol. 44, no. 1, pp. 34–47, 2020, doi: 10.3906/biy-1908-32.

[36] N. Kalaji, A. Deloge, N. Sheibat-Othman, O. Boyron, I. About, and H. Fessi, “Controlled Release Carriers of Growth Factors FGF-2 and TGF 1: Synthesis, Characterization and Kinetic Modelling,” (in English), Journal of Biomedical Nanotechnology, vol. 6, no. 2, pp. 106–116, 2010-04-01 2010, doi: 10.1166/jbn.2010.1102.

[37] X. J. Loh, V. P. Nam Nguyen, N. Kuo, and J. Li, “Encapsulation of basic fibroblast growth factor in thermogelling copolymers preserves its bioactivity,” Journal of Materials Chemistry, vol. 21, no. 7, pp. 2246–2254, 2011, doi: 10.1039/c0jm03051a.

[38] M. J. Hernandez et al., “Decellularized Extracellular Matrix Hydrogels as a Delivery Platform for MicroRNA and Extracellular Vesicle Therapeutics,” (in eng), Adv Ther (Weinh), vol. 1, no. 3, Jul 2018, doi: 10.1002/adtp.201800032.

[39] R. Bhatta et al., “Injectable extracellular vesicle hydrogels with tunable viscoelasticity for depot vaccine,” Nature Communications, vol. 16, no. 1, p. 3781, 2025/04/22 2025, doi: 10.1038/s41467-025-59278-0.

[40] A. Hashemi, M. Ezati, M. P. Nasr, I. Zumberg, and V. Provaznik, “Extracellular Vesicles and Hydrogels: An Innovative Approach to Tissue Regeneration,” ACS Omega, vol. 9, no. 6, pp. 6184–6218, 2024, doi: 10.1021/acsomega.3c08280.

[41] M. Ernits et al., “Microfluidic production, stability and loading of synthetic giant unilamellar vesicles,” Scientific Reports, vol. 14, no. 1, p. 14071, 2024/06/18 2024, doi: 10.1038/s41598-024-64613-4.

[42] O. Staufer et al., “Microfluidic production and characterization of biofunctionalized giant unilamellar vesicles for targeted intracellular cargo delivery,” (in eng), Biomaterials, vol. 264, p. 120203, Jan 2021, doi: 10.1016/j.biomaterials.2020.120203.

[43] A. Llopis-Lorente, M. J. G. Schotman, H. V. Humeniuk, J. C. M. van Hest, P. Y. W. Dankers, and L. K. E. A. Abdelmohsen, “Artificial cells with viscoadaptive behavior based on hydrogel-loaded giant unilamellar vesicles,” Chemical Science, vol. 15, no. 2, pp. 629–638, 2024, doi: 10.1039/d3sc04687g.

[44] S. W. Hwang et al., “Hybrid Vesicles Enable Mechano-Responsive Hydrogel Degradation,” (in eng), Angew Chem Int Ed Engl, vol. 62, no. 41, p. e202308509, Oct 9 2023, doi: 10.1002/anie.202308509.

[45] N. Ashammakhi et al., “Bioinks and bioprinting technologies to make heterogeneous and biomimetic tissue constructs,” (in eng), Mater Today Bio, vol. 1, p. 100008, Jan 2019, doi: 10.1016/j.mtbio.2019.100008.

[46] A. Schwab, R. Levato, M. D’Este, S. Piluso, D. Eglin, and J. Malda, “Printability and Shape Fidelity of Bioinks in 3D Bioprinting,” Chemical Reviews, vol. 120, no. 19, pp. 10850–10877, 2020, doi: 10.1021/acs.chemrev.0c00084.

[47] J. V. Jester, D. Brown, A. Pappa, and V. Vasiliou, “Myofibroblast differentiation modulates keratocyte crystallin protein expression, concentration, and cellular light scattering,” (in eng), Invest Ophthalmol Vis Sci, vol. 53, no. 2, pp. 770–8, Feb 16 2012, doi: 10.1167/iovs.11-9092.

[48] J. W. Foster, R. M. Gouveia, and C. J. Connon, “Low-glucose enhances keratocyte-characteristic phenotype from corneal stromal cells in serum-free conditions,” (in eng), Sci Rep, vol. 5, p. 10839, Jun 3 2015, doi: 10.1038/srep10839.

[49] V. Jhanji, J. Mehta, M. Funderburgh, A. Riau, and G. Yam, “Keratocyte biology,” Experimental Eye Research, vol. 196, p. 108062, 2020/07/01 2020, doi: 10.1016/j.exer.2020.108062.

[50] G. H.-F. Yam et al., “Ex Vivo Propagation of Human Corneal Stromal “Activated Keratocytes” for Tissue Engineering,” Cell Transplantation, vol. 24, no. 9, pp. 1845–1861, 2015, doi: 10.3727/096368914x685069.

[51] S. E. Wilson, “Corneal myofibroblasts and fibrosis,” (in eng), Exp Eye Res, vol. 201, p. 108272, Dec 2020, doi: 10.1016/j.exer.2020.108272.

[52] K. Poole, K. S. Iyer, D. W. Schmidtke, W. M. Petroll, and V. D. Varner, “Corneal Keratocytes, Fibroblasts, and Myofibroblasts Exhibit Distinct Transcriptional Profiles In Vitro,” (in eng), Invest Ophthalmol Vis Sci, vol. 66, no. 3, p. 28, Mar 3 2025, doi: 10.1167/iovs.66.3.28.

[53] E. Hermann, S. Bleicken, Y. Subburaj, and A. J. García-Sáez, “Automated analysis of giant unilamellar vesicles using circular Hough transformation,” Bioinformatics, vol. 30, no. 12, pp. 1747–1754, 2014, doi: 10.1093/bioinformatics/btu102.

[54] F. Dehli et al., “Biobased photocrosslinkable gelatin-methacrylate hydrogels promote the growth and phenotype maintenance of human corneal keratocytes,” Materials Advances, vol. 6, no. 12, pp. 3805–3816, 2025, doi: 10.1039/d5ma00076a.

[55] M. R. Birajdar, K. K. Kaushik, S. Sharma, R. K. Sah, A. Archana, and S. Tiwari, “Discharge behavior of liposomal vesicles embedded in gellan gum matrix: The effect of plasticizers,” Colloids and Surfaces A: Physicochemical and Engineering Aspects, vol. 716, p. 136660, 2025/07/05/ 2025, doi: 10.1016/j.colsurfa.2025.136660.

[56] Y. Zheng, C. Pan, P. Xu, and K. Liu, “Hydrogel-mediated extracellular vesicles for enhanced wound healing: the latest progress, and their prospects for 3D bioprinting,” Journal of Nanobiotechnology, vol. 22, no. 1, pp. 57–57, 2024/12// 2024, doi: 10.1186/S12951-024-02315-9.

[57] C. Kilic Bektas and V. Hasirci, “Cell loaded 3D bioprinted GelMA hydrogels for corneal stroma engineering,” (in eng), Biomater Sci, vol. 8, no. 1, pp. 438–449, Dec 17 2019, doi: 10.1039/c9bm01236b.

[58] R. Chand, G. Janarthanan, K. Elkhoury, and S. Vijayavenkataraman, “Digital light processing 3D bioprinting of biomimetic corneal stroma equivalent using gelatin methacryloyl and oxidized carboxymethylcellulose interpenetrating network hydrogel,” (in eng), Biofabrication, vol. 17, no. 2, Feb 10 2025, doi: 10.1088/1758-5090/adab27.

[59] D. F. Duarte Campos et al., “Corneal bioprinting utilizing collagen-based bioinks and primary human keratocytes,” Journal of Biomedical Materials Research Part A, vol. 107, no. 9, pp. 1945–1953, 2019, doi: 10.1002/jbm.a.36702.

[60] A. Ghosh, A. K. Bera, V. Singh, S. Basu, and F. Pati, “Bioprinting of anisotropic functional corneal stroma using mechanically robust multi-material bioink based on decellularized cornea matrix,” (in eng), Biomater Adv, vol. 165, p. 214007, Dec 2024, doi: 10.1016/j.bioadv.2024.214007.

[61] H. Kim, M. N. Park, J. Kim, J. Jang, H. K. Kim, and D. W. Cho, “Characterization of cornea-specific bioink: high transparency, improved in vivo safety,” (in eng), J Tissue Eng, vol. 10, p. 2041731418823382, Jan-Dec 2019, doi: 10.1177/2041731418823382.

[62] T. Kawakita et al., “Preservation and expansion of the primate keratocyte phenotype by downregulating TGF-beta signaling in a low-calcium, serum-free medium,” (in eng), Invest Ophthalmol Vis Sci, vol. 47, no. 5, pp. 1918–27, May 2006, doi: 10.1167/iovs.05-1040.

[63] M. L. Funderburgh, M. M. Mann, and J. L. Funderburgh, “Keratocyte phenotype is enhanced in the absence of attachment to the substratum,” (in eng), Mol Vis, vol. 14, pp. 308–17, Feb 9 2008.

[64] D. T. Chu et al., “An Update on the Progress of Isolation, Culture, Storage, and Clinical Application of Human Bone Marrow Mesenchymal Stem/Stromal Cells,” (in eng), Int J Mol Sci, vol. 21, no. 3, Jan 21 2020, doi: 10.3390/ijms21030708.

[65] K. Bieback, K. Schallmoser, H. Klüter, and D. Strunk, “Clinical Protocols for the Isolation and Expansion of Mesenchymal Stromal Cells,” (in eng), Transfus Med Hemother, vol. 35, no. 4, pp. 286–294, 2008, doi: 10.1159/000141567.

[66] S. Bhat, P. Viswanathan, S. Chandanala, S. J. Prasanna, and R. N. Seetharam, “Expansion and characterization of bone marrow derived human mesenchymal stromal cells in serum-free conditions,” (in eng), Sci Rep, vol. 11, no. 1, p. 3403, Feb 9 2021, doi: 10.1038/s41598-021-83088-1.

