## Supplementary Information for "Engineered cell-like vesicles instruct keratocyte differentiation for corneal biofabrication and regeneration"

This supplementary information contains:

- Figure S1
- Table S1
- Figure S2
- Figure S3
- Figure S4
- Figure S5
- Figure S6
- Figure S7

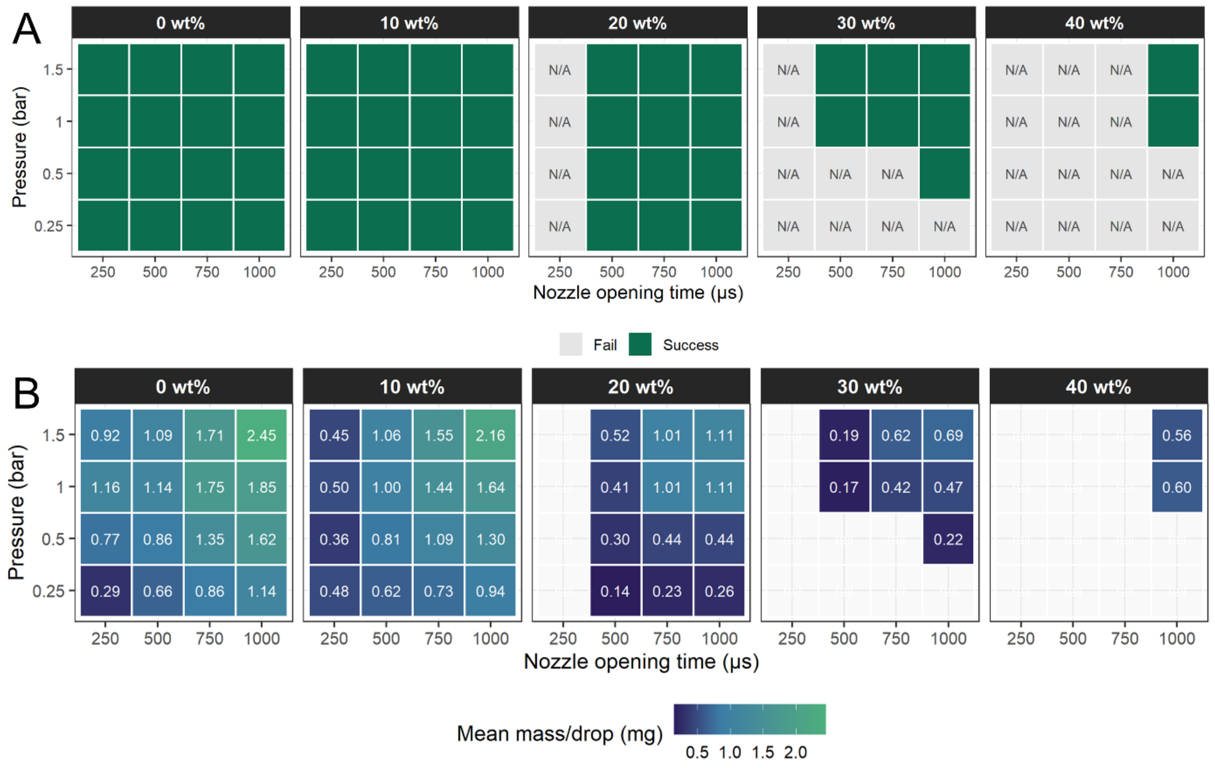


**Figure S1. Printability and droplet mass analysis of RA-GelMA droplets.** (A) Printability map of RA-GelMA formulations at 0, 10, 20, 30, and 40 wt % across different printing pressures and nozzle opening times. Green tiles indicate successful printing, while grey tiles indicate non-printable conditions. (B) Mean droplet mass obtained under each printing condition. Values indicate the mean mass per droplet in milligrams, calculated from the collection of 10 droplets per condition. Non-printable conditions are shown as failed conditions.

**Table S1.** Mean droplet mass across RA-GelMA concentrations and printing conditions

| Concentration | Pressure | 250 µs | 500 µs | 750 µs | 1000 µs |
|---|---|---|---|---|---|
| 0 wt % | 0.25 bar | 0.29 ± 0.03 | 0.66 ± 0.03 | 0.86 ± 0.03 | 1.14 ± 0.03 |
| 0 wt % | 0.5 bar | 0.77 ± 0.02 | 0.86 ± 0.02 | 1.35 ± 0.01 | 1.62 ± 0.01 |
| 0 wt % | 1.0 bar | 1.16 ± 0.02 | 1.14 ± 0.02 | 1.75 ± 0.04 | 1.85 ± 0.03 |
| 0 wt % | 1.5 bar | 0.92 ± 0.02 | 1.09 ± 0.09 | 1.71 ± 0.07 | 2.45 ± 0.04 |
| 10 wt % | 0.25 bar | 0.48 ± 0.08 | 0.62 ± 0.07 | 0.73 ± 0.04 | 0.94 ± 0.02 |
| 10 wt % | 0.5 bar | 0.36 ± 0.01 | 0.81 ± 0.02 | 1.09 ± 0.02 | 1.30 ± 0.02 |
| 10 wt % | 1.0 bar | 0.50 ± 0.06 | 1.00 ± 0.02 | 1.44 ± 0.03 | 1.64 ± 0.03 |
| 10 wt % | 1.5 bar | 0.45 ± 0.02 | 1.06 ± 0.01 | 1.55 ± 0.02 | 2.16 ± 0.24 |
| 20 wt % | 0.25 bar | N/A | 0.14 ± 0.06 | 0.23 ± 0.01 | 0.26 ± 0.01 |
| 20 wt % | 0.5 bar | N/A | 0.30 ± 0.04 | 0.44 ± 0.02 | 0.44 ± 0.02 |
| 20 wt % | 1.0 bar | N/A | 0.41 ± 0.03 | 1.01 ± 0.08 | 1.11 ± 0.09 |
| 20 wt % | 1.5 bar | N/A | 0.52 ± 0.08 | 1.01 ± 0.02 | 1.11 ± 0.01 |
| 30 wt % | 0.25 bar | N/A | N/A | N/A | N/A |
| 30 wt % | 0.5 bar | N/A | N/A | N/A | 0.22 ± 0.03 |
| 30 wt % | 1.0 bar | N/A | 0.17 ± 0.01 | 0.42 ± 0.07 | 0.47 ± 0.04 |
| 30 wt % | 1.5 bar | N/A | 0.19 ± 0.02 | 0.62 ± 0.05 | 0.69 ± 0.11 |
| 40 wt % | 0.25 bar | N/A | N/A | N/A | N/A |
| 40 wt % | 0.5 bar | N/A | N/A | N/A | N/A |
| 40 wt % | 1.0 bar | N/A | N/A | N/A | 0.60 ± 0.10 |
| 40 wt % | 1.5 bar | N/A | N/A | N/A | 0.56 ± 0.05 |

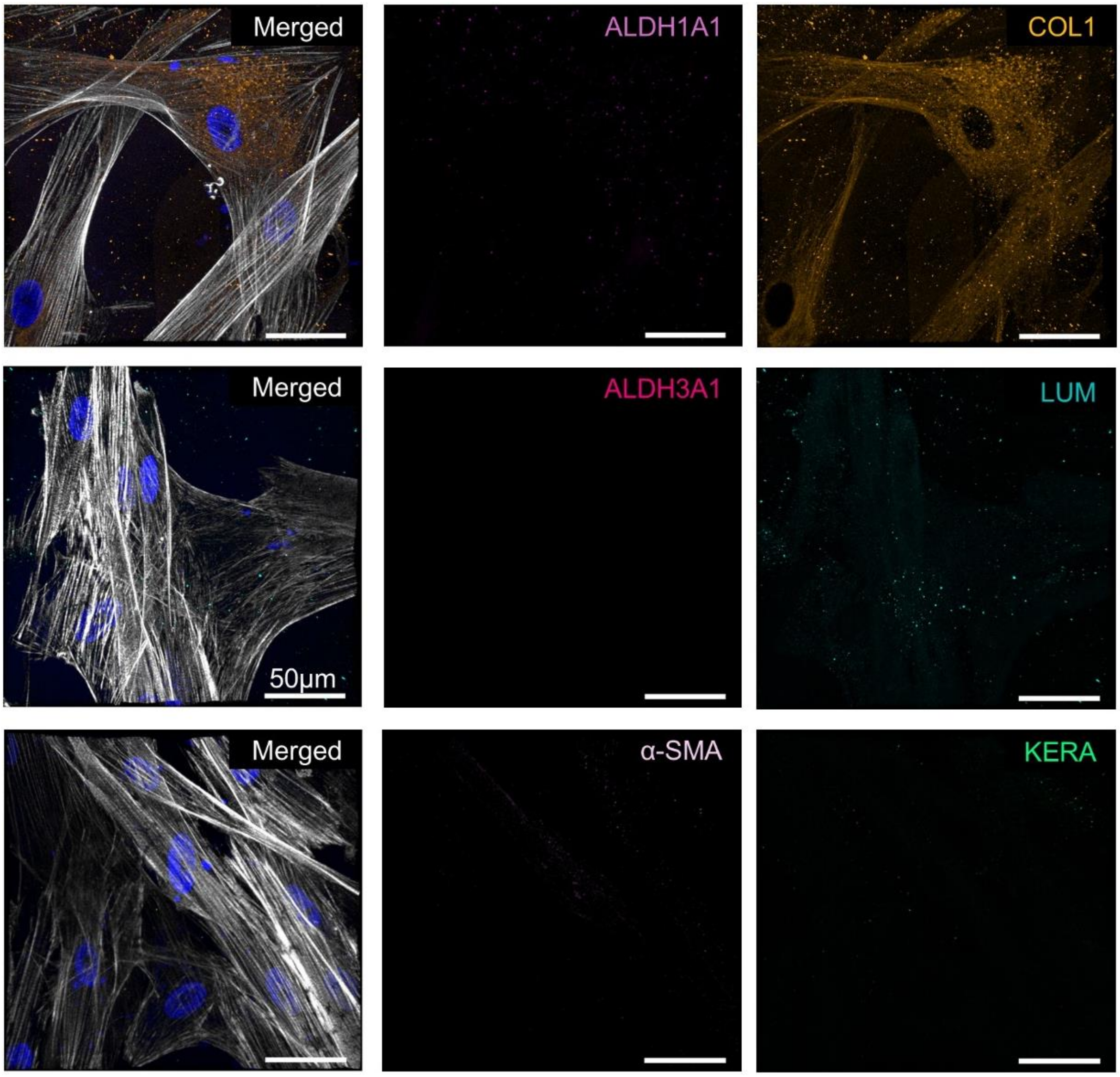


**Figure S2. Absence of keratocyte marker expression in BM-MSC cultured under basal conditions.** BM-MSC ($2 \times 10^6$ cells $mL^{-1}$) were encapsulated in 30 wt% RA-GelMA and cultured for 17 days without differentiation factors. Constructs were stained for ALDH1A1 (light pink), ALDH3A1 (dark pink), Lumican (turquoise), Keratocan (green), Collagen I (orange), and α-SMA (lilac). Actin filaments were stained with phalloidin (white) and nuclei with Hoechst (blue). Scale bars = 50 µm.

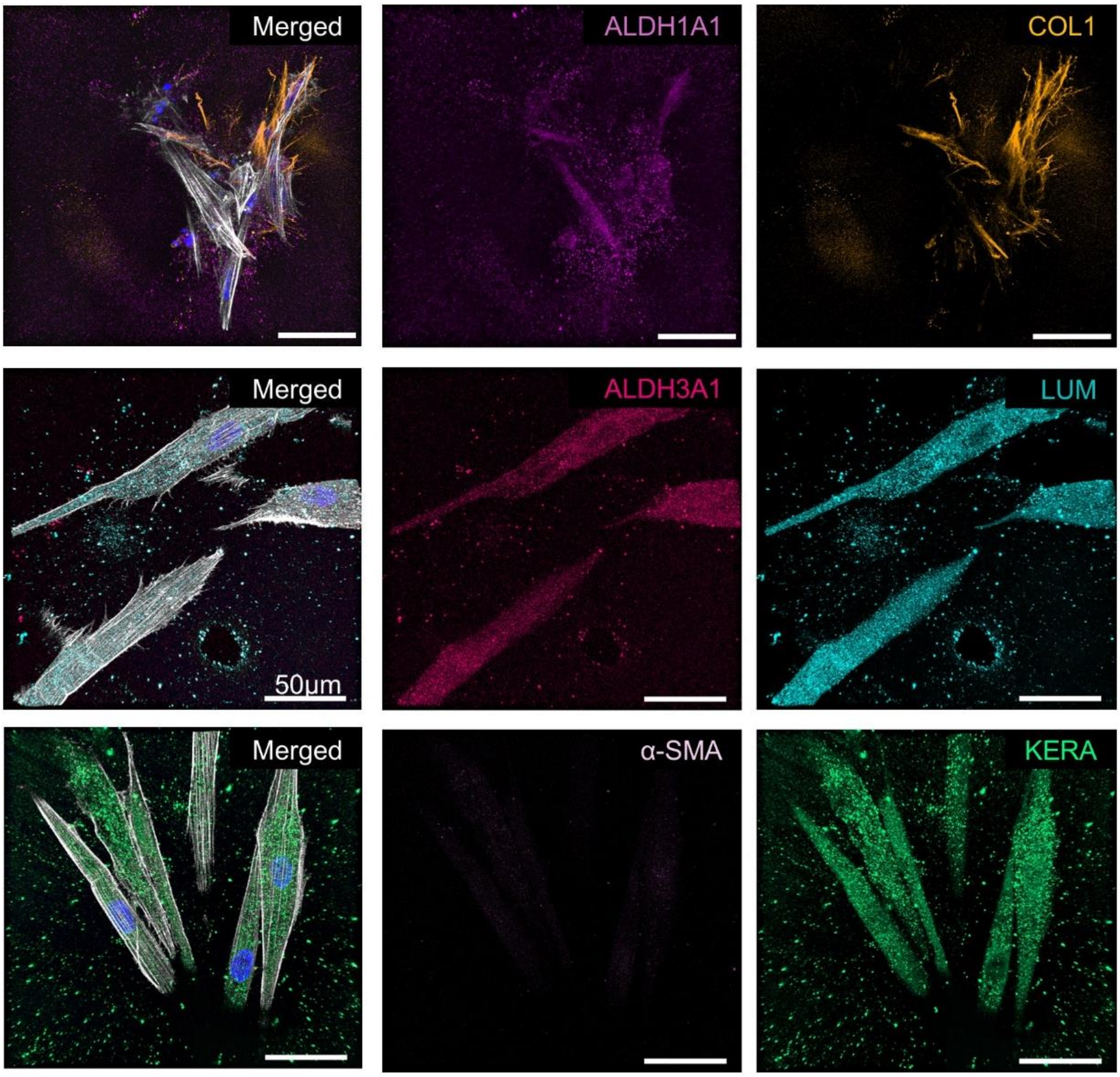


**Figure S3. Immunofluorescence analysis of keratocyte differentiation induced by soluble growth factor supplementation.** BM-MSC ($2 \times 10^6$ cells $mL^{-1}$) were encapsulated in 30 wt% RA-GelMA and cultured for 3 days prior to induction of keratocyte differentiation. Constructs were subsequently cultured for 14 days in keratocyte differentiation medium supplemented with soluble FGF-2 and TGF-β3. Cells were stained for ALDH1A1 (light pink), ALDH3A1 (dark pink), Lumican (turquoise), Keratocan (green), Collagen I (orange), and α-SMA (lilac). Actin filaments were stained with phalloidin (white) and nuclei with Hoechst (blue). Scale bars = 50 µm.

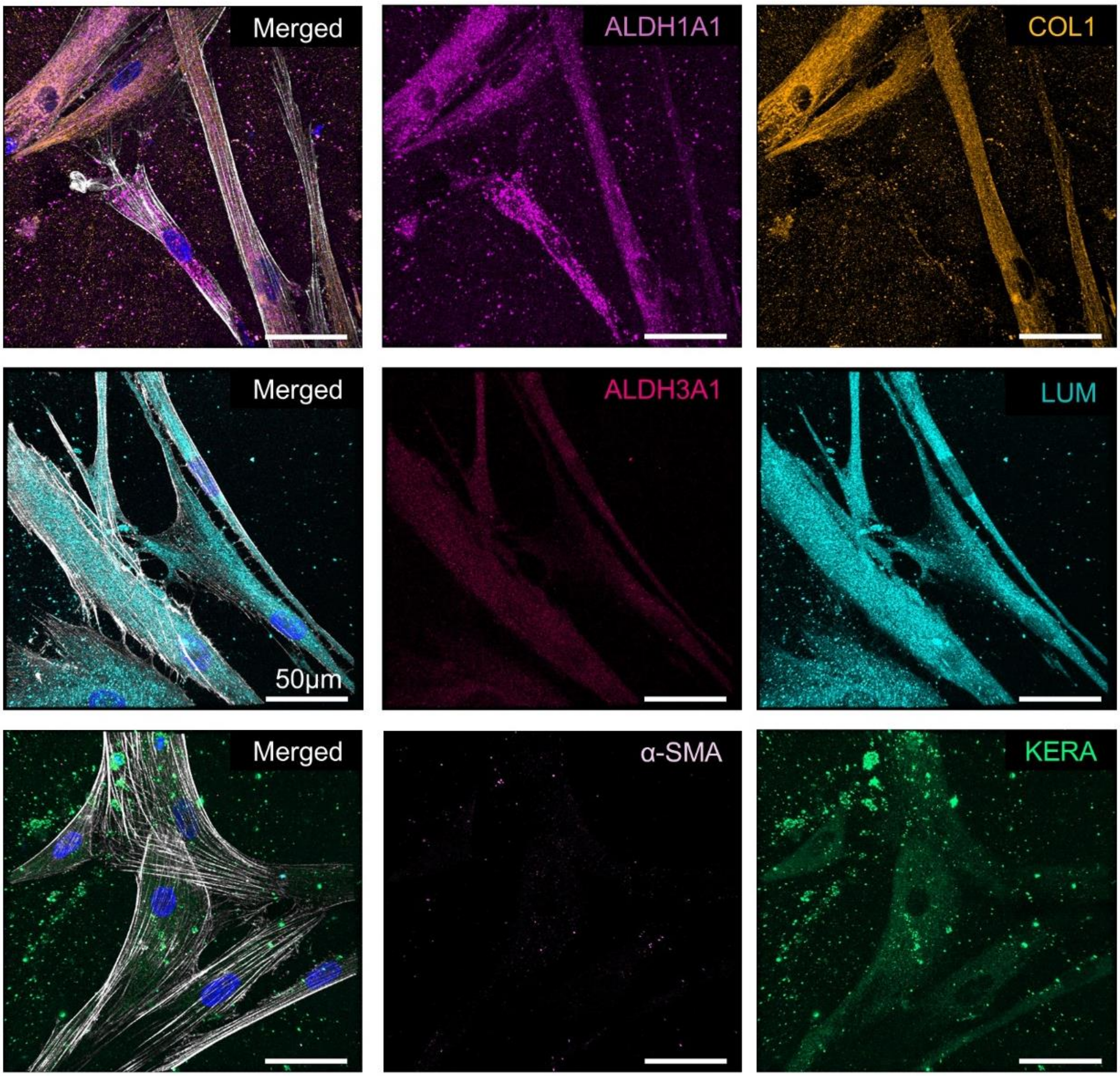


**Figure S4. Immunofluorescence analysis of keratocyte differentiation induced by GUVs-mediated growth factor delivery.** BM-MSC ($2 \times 10^6$ cells $mL^{-1}$) were encapsulated in 30 wt% RA-GelMA together with GUVs co-loaded with FGF-2 and TGF-β3. After 3 days of culture, constructs were maintained for 14 days in growth factor-free keratocyte differentiation medium. Cells were stained for ALDH1A1 (light pink), ALDH3A1 (dark pink), Lumican (turquoise), Keratocan (green), Collagen I (orange), and α-SMA (lilac). Actin filaments were stained with phalloidin (white) and nuclei with Hoechst (blue). Scale bars = 50 μm.

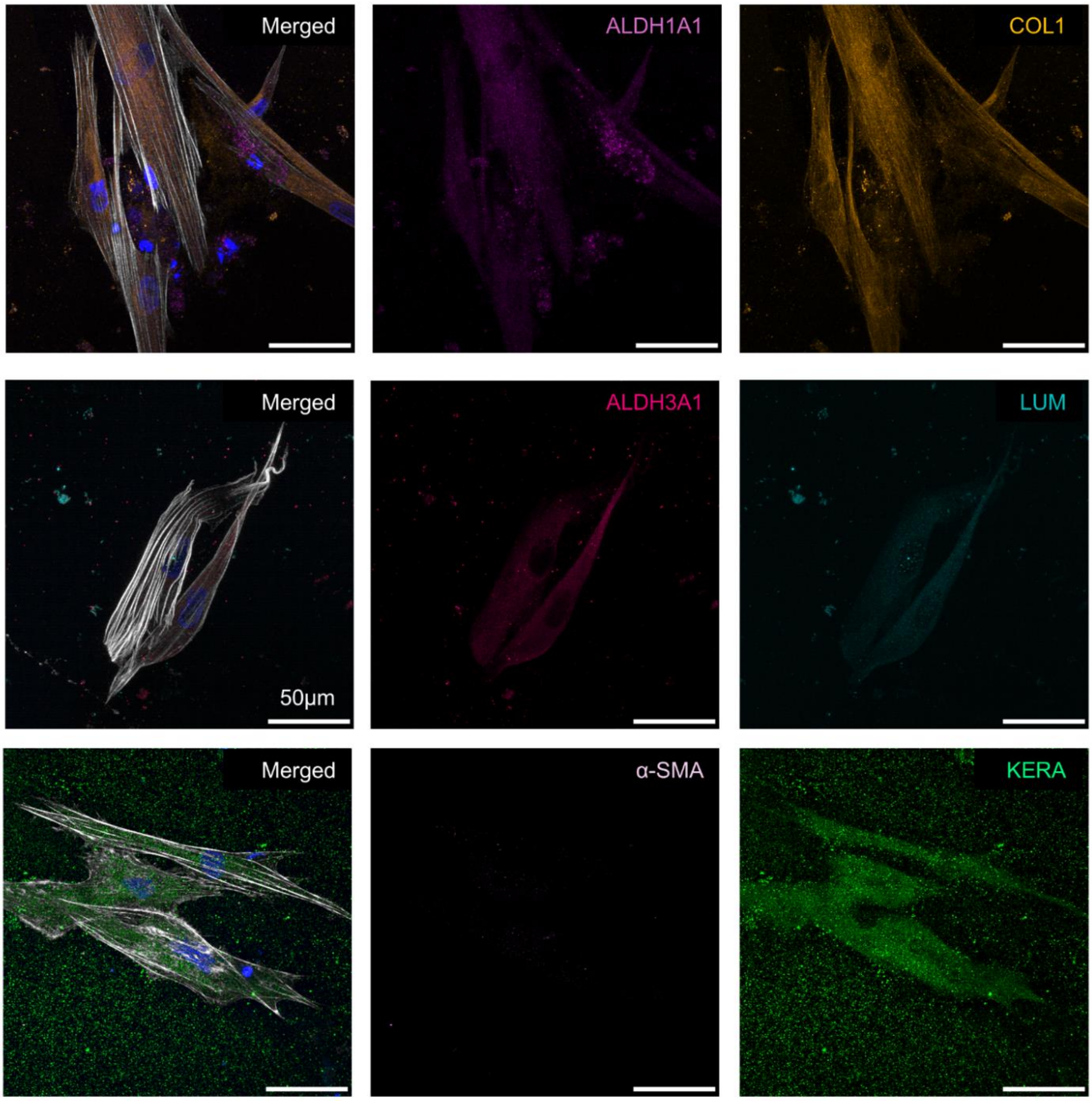


**Figure S5. Immunofluorescence analysis of keratocyte differentiation following drop-on-demand bioprinting of BM-MSC-laden RA-GelMA bioinks containing growth factor-loaded GUVs.** BM-MSC ($2 \times 10^6$ cells $mL^{-1}$) were bioprinted in 30 wt% RA-GelMA together with GUVs co-loaded with FGF-2 and TGF-β3. Following 3 days of culture, a 14-day keratocyte differentiation period was initiated in growth factor-free keratocyte differentiation medium. Constructs were stained for ALDH1A1 (light pink), ALDH3A1 (dark pink), Lumican (turquoise), Keratocan (green), Collagen I (orange), and α-SMA (lilac). Actin filaments were stained with phalloidin (white) and nuclei with Hoechst (blue). Scale bars = 50 µm.

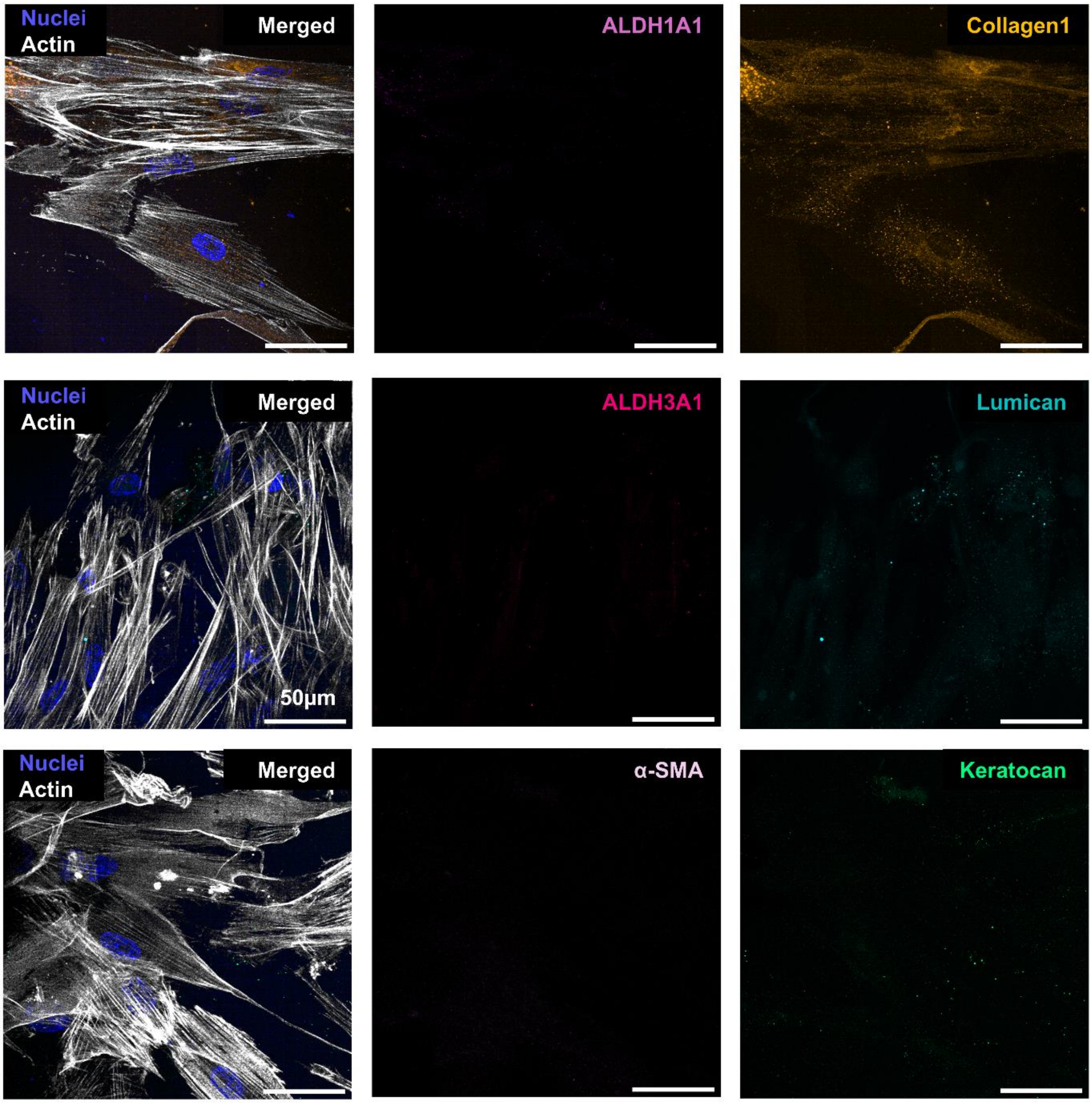


**Figure S6. Absence of keratocyte marker expression in BM-MSC printed and maintained under basal conditions.** BM-MSC ($2 \times 10^6$ cells $mL^{-1}$) were encapsulated in 30 wt% RA-GelMA and cultured for 17 days without differentiation factors. Constructs were stained for ALDH1A1 (light pink), ALDH3A1 (dark pink), Lumican (turquoise), Keratocan (green), Collagen I (orange), and α-SMA (lilac). Actin filaments were stained with phalloidin (white) and nuclei with Hoechst (blue). Scale bars = 50 µm.

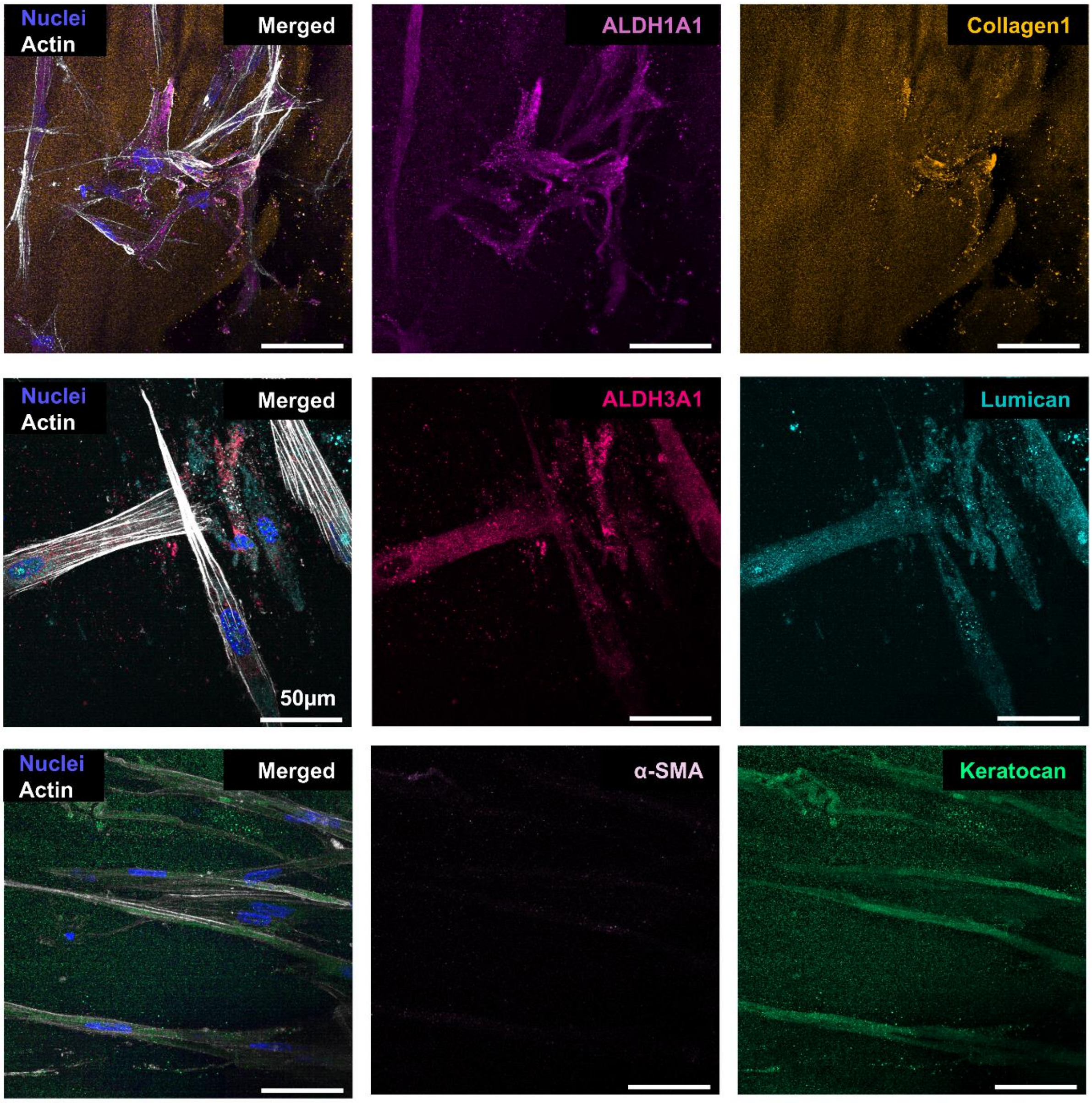


**Figure S7. Immunofluorescence analysis of keratocyte differentiation induced by soluble growth factor supplementation post printing.** BM-MSC ($2 \times 10^6$ cells $mL^{-1}$) were printed in 30 wt% RA-GelMA and cultured for 3 days prior to induction of keratocyte differentiation. Constructs were subsequently cultured for 14 days in keratocyte differentiation medium supplemented with soluble FGF-2 and TGF-β3. Cells were stained for ALDH1A1 (light pink), ALDH3A1 (dark pink), Lumican (turquoise), Keratocan (green), Collagen I (orange), and α-SMA (lilac). Actin filaments were stained with phalloidin (white) and nuclei with Hoechst (blue). Scale bars = 50 µm.
